# NETWORKED proteins link actin-dependent vacuolar organization to the endoplasmic reticulum

**DOI:** 10.1101/2023.09.29.560113

**Authors:** Sabrina Kaiser, Dietmar Mehlhorn, Paulina Ramirez Miranda, Ana Carolina Vilchez, Fabian Ries, Nadine Field, Frederik Sommer, Michael Schroda, Karin Schumacher, Verena Kriechbaumer, Felix Willmund, Till Ischebeck, Christopher Grefen, David Scheuring

## Abstract

The organization of plant endomembrane compartments depends on coordinated interactions between membranes and the actin cytoskeleton. NETWORKED (NET) proteins are plant-specific, membrane-associated actin-binding proteins whose cellular functions remain incompletely understood. Here, we show that NET3 and NET4 have overlapping functions in maintaining vacuolar architecture and that NET4 connects actin organization with vacuolar remodeling. NET4A overexpression altered actin filament organization, partially rescued vacuolar defects in the *act7-4* mutant, and promoted recovery of vacuolar continuity following actin depolymerization and latrunculin B washout. NET proteins also associated with RABG3 GTPases involved in vacuolar organization. In addition, NET4A and NET4B localized to the endoplasmic reticulum (ER), associated with the ER membrane contact site protein VAP27-1, and NET4B interacted with the lipid-transfer protein ORP2A. Together, these findings raise the possibility that NET3 and NET4 proteins contribute to communication between the ER and the vacuole.

## Introduction

Compartmentalization of cells is a hallmark of all eukaryotes. Exchange of material and communication between organelles are secured by vesicular transport and by the formation of membrane contact sites (MCSs). Protein transport requires sorting into vesicles via coat proteins, interaction with GTPases, tethering and eventual SNARE (soluble N-ethylmaleimide-sensitive factor attachment receptor)-mediated fusion with the target membrane. In order to target proteins to the lytic vacuole, different trafficking routes can be taken (Cui et al., 2020). The best studied pathway towards the vacuole includes the sequential recruitment of Rab5-like and Rab7-like GTPases, which act as molecular switches by cycling between inactive GDP-bound and active GTP-bound states (Ebine et al., 2014). Notably, Rab7-like proteins, called RABGs in plants, interact with the HOPS (HOMOTYPIC FUSION AND VACUOLE PROTEIN SORTING) tethering complex at the vacuole (Takemoto et al., 2018). Together with the CORVET (CLASS C CORE VACUOLE/ENDOSOME TETHERING) complex, HOPS belongs to the class C Vps tethering complexes that coordinate endosomal and vacuolar trafficking. HOPS and CORVET share the core subunits VPS16, VPS18, VPS11 and VPS33, whereas HOPS additionally contains VPS39 and VPS41, which confer specificity for Rab7/Ypt7-positive late endosomal and vacuolar membranes (Balderhaar and Ungermann, 2013; Aniento et al., 2022). In yeast, recruitment of HOPS by the Rab7 homolog Ypt7 is central to vacuolar membrane tethering and SNARE-dependent homotypic vacuole fusion, illustrating how Rab GTPases, tethering complexes and membrane fusion are functionally integrated (Balderhaar and Ungermann, 2013; Shvarev et al., 2022). In plants, impairment of HOPS components has severe developmental consequences, while induced interference with HOPS function results in dramatic vacuolar fragmentation (Brillada et al., 2018; Takemoto et al., 2018).

MCSs provide an additional mechanism for communication between organelles. These regions bring membranes into close proximity and can coordinate lipid exchange, signaling, membrane remodeling and organelle dynamics (Scorrano et al., 2019). In yeast and animals, numerous contact sites have been characterized, with the endoplasmic reticulum (ER) serving as a central component of many of these structures. In plants, investigations of MCSs have come more recently into focus, and the best characterized examples include ER–plasma membrane contact sites (EPCS), whose formation involves a complex interplay between actin filaments, actin-recruiting proteins, microtubules and integral ER proteins (Wang et al., 2014; Zang et al., 2021). More recently, ER–chloroplast and ER–autophagosome contact sites have also been described (Renna et al., 2024; Ye et al., 2022).

A particularly relevant example from yeast is the nucleus–vacuole junction (NVJ), a specialized ER–vacuole membrane contact site formed by the interaction of the ER-localized protein Nvj1 with the vacuolar membrane protein Vac8 (Pan et al., 2000). The NVJ represents a specialized membrane domain involved in organelle communication, membrane remodeling and lipid metabolism. Lipid-associated proteins are enriched at the NVJ, and recent work has connected ER–vacuole contact sites with sphingolipid metabolism and the regulation of vacuolar morphology. Vacuolar membranes themselves contain distinct lipid domains that are associated with membrane trafficking, organelle contacts and homotypic fusion, further highlighting the close relationship between membrane composition and vacuolar architecture (Hurst and Fratti, 2020). While functionally equivalent ER–vacuole membrane contact sites in plants are only now emerging, the ER is already known to be intimately connected to vacuole biogenesis, serving as an essential membrane source for lytic vacuole formation (Viotti et al., 2013; Scheuring et al., 2024). Together, these observations suggest that ER–vacuole membrane organization and lipid homeostasis may be closely linked to the development and maintenance of vacuolar architecture.

In plants, members of the NETWORKED (NET) family possess actin-binding capacity and localize to different membrane compartments. NET proteins were identified as a plant-specific superfamily of actin-binding proteins that simultaneously associate with the actin cytoskeleton and specific membrane compartments, suggesting that they provide molecular interfaces between actin and membranes (Deeks et al., 2012). Different NET proteins link actin to specific membranes, including the tonoplast, nuclear membrane, plasma membrane, plasmodesmata and ER (Deeks et al., 2012; Wang et al., 2014; Kaiser et al., 2019; Zang et al., 2021; Hawkins et al., 2023). Actin filaments and microtubules represent the molecular tracks for moving organelles and molecules within a cell (Vale, 2003). While in animals microtubules have major functions in vesicle trafficking and organelle organization, actin filaments play a particularly prominent role in these processes in plants (Kost and Chua, 2002). The connection between actin and the vacuole is mediated in part by the NET4 protein family. NET4A localizes to the tonoplast and to highly constricted regions of the vacuole, and its abundance influences vacuolar morphology (Kaiser et al., 2019). Whereas the *net4a net4b* double mutant showed more spherical and larger vacuoles, overexpression of NET4A rendered vacuoles more spherical, smaller and more condensed around the nucleus (Kaiser et al., 2019). These changes in vacuolar morphology are closely connected to plant growth. We previously showed that actin-dependent vacuolar occupancy contributes to auxin-induced growth repression (Scheuring et al., 2016), while subsequent work demonstrated that vacuolar occupancy is a general determinant of cell elongation and growth (Kaiser et al., 2021). Thus, the organization of the actin–vacuole interface represents an important component of plant cell expansion.

The molecular relationship between NET4, actin and the tonoplast has recently been further defined. Hawkins et al. (2023) demonstrated that NET4A and NET4B interact with F-actin and with active RABG3 GTPases through distinct domains, allowing simultaneous association with actin filaments and the tonoplast. The NET4–RABG3 module contributes to cytoskeletal remodeling during stomatal immunity, establishing a direct molecular connection between actin organization and RABG3-dependent vacuolar membrane regulation. NET proteins therefore provide an attractive molecular framework through which cytoskeletal organization can be coupled to membrane identity and dynamics. NET proteins are also associated with ER membrane organization. NET3C interacts with the ER-integral membrane protein VAP27 and contributes to the organization of the actin and microtubule cytoskeletons at ER–plasma membrane contact sites (Wang et al., 2014). The subsequent identification of a NET3C–KLCR1–IQD2 complex further demonstrated that NET3C participates in the organization of the cytoskeleton and ER at ER–PM contact sites (Zang et al., 2021). These findings indicate that NET proteins can function as molecular organizers that couple actin to membrane-associated protein complexes and specialized membrane domains. Despite these advances, it remains unclear how different NET proteins cooperate to organize the vacuolar system and how actin-dependent vacuolar organization is connected to ER-associated membrane processes and lipid homeostasis. In particular, the functional relationship between NET3 and NET4 proteins has remained largely unexplored, although both contain conserved domains involved in membrane and actin association. Moreover, whether NET proteins connect RABG3-dependent vacuolar organization with ER-associated membrane domains and lipid-associated proteins remains unknown.

Here, we investigated the overlapping functions of NET3 and NET4 in vacuolar organization and their relationship to the actin cytoskeleton. We found that NET4A abundance influences actin organization and vacuolar architecture, partially rescues vacuolar defects in the *act7-4* mutant, and promotes recovery of vacuolar continuity following actin depolymerization. Our analyses further revealed associations between NET3 and NET4 proteins and RABG3 GTPases, as well as genetic interactions between NET4 and the RABG3- and HOPS-dependent machinery involved in vacuolar organization. Beyond these connections to vacuolar organization, NET4 proteins localize to the endoplasmic reticulum (ER), interact with the ER membrane protein VAP27-1, and NET4B associates with the lipid-transfer protein ORP2A. These findings connect NET3 and NET4 proteins to vacuolar organization and ER-associated proteins, providing a basis for investigating whether they contribute to communication between the ER and the vacuole.

## Results

### NET3 and NET4 proteins redundantly regulate vacuolar architecture

Previous analyses of NET4A and NET4B showed that changes in NET4 protein abundance affect vacuolar morphology, whereas individual *net4* mutants display relatively mild phenotypes (Kaiser 2019, Hawkins 2023). Since NET3A and NET3C, like NET4A and NET4B, contain the conserved IRQ domain involved in tonoplast-localized RABG3 binding, we asked whether NET3 proteins contribute to vacuolar organization together with NET4. We generated single CRISPR/Cas9-based deletion mutants as well as higher-order mutant combinations by either deleting several genes via CRISPR/Cas9 or combining T-DNA insertion alleles with CRISPR/Cas9-mediated gene deletions (Fig. S1) and analyzed vacuolar morphology using the vacuolar morphology index, VMI (Löfke et al., 2015).

Pronounced defects in vacuolar organization were observed in *net3c net4a net4b* triple mutants but not a *net3c* single mutant (Fig. 1A). Vacuoles in these plants were less compact and occupied a smaller fraction of the cell, often accompanied with small, isolated vacuolar subdomains (Fig. 1B). In contrast, *net3a net4a net4b* mutants did not show a comparable reduction in VMI or occupancy (Fig. S2). Notably, the vacuolar phenotype of the *net3c net4a net4b* mutant was accompanied by reduced root growth (Fig. 1C) which could be rescued by reintroduction of NET4B (Fig. 1D). This points towards a functional overlap between NET3 and NET4 proteins in the regulation of vacuolar architecture. The relatively mild phenotypes of the lower-order mutants indicate that NET3 and NET4 can compensate for one another, whereas simultaneous loss of NET3C and NET4A/B results in pronounced defects in vacuolar morphology and root growth. Consistent with overlapping functions among NET family members, overexpression of NET4B similarly affected vacuolar occupancy and root growth (Fig. S3).

**Fig. 1.**
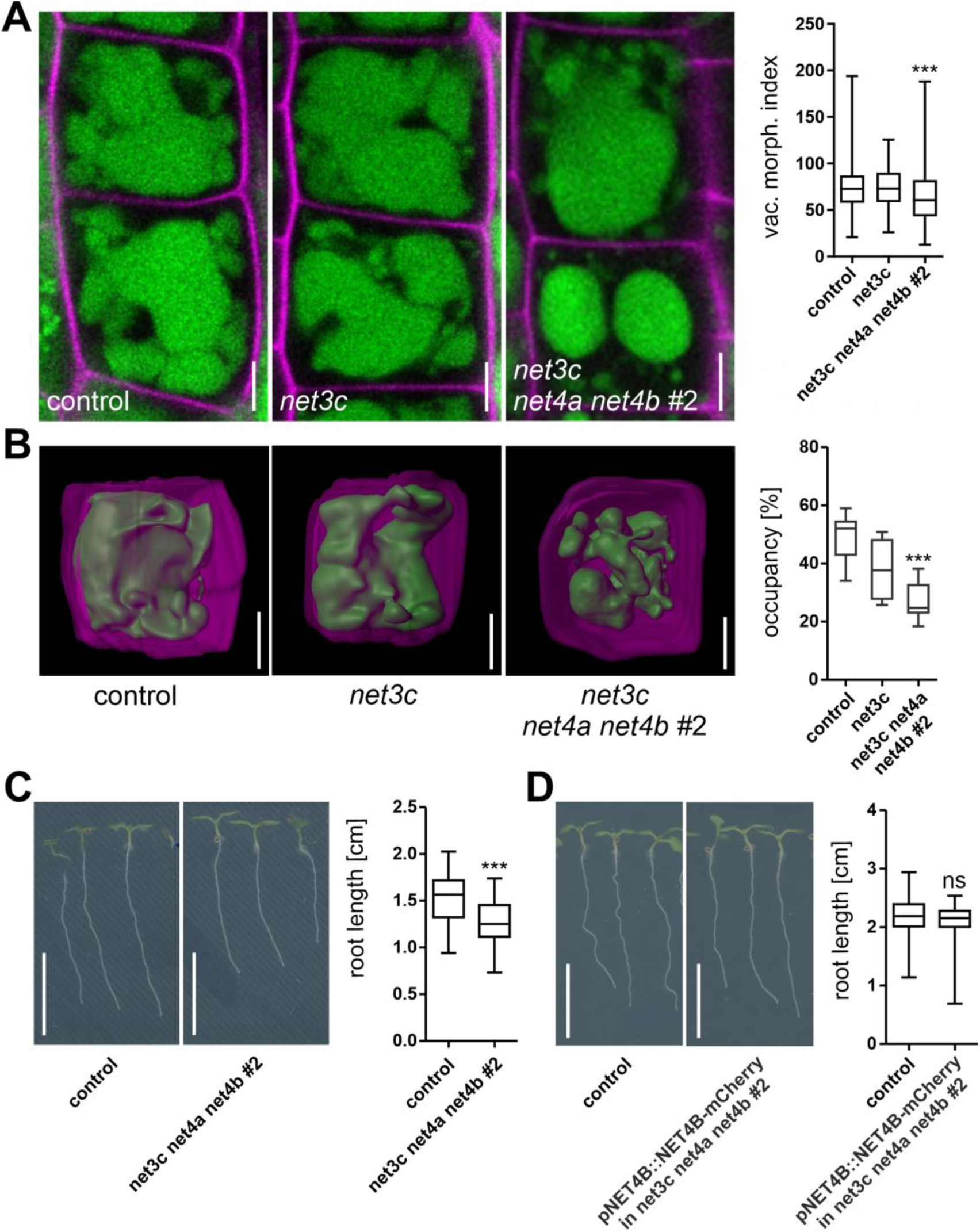
Phenotypic analysis of NET3–NET4 multiple mutants. **(A)** Vacuolar morphology in the CRISPR/Cas9-derived *net3c* mutant (*n* = 84) and the *net3c net4a net4b #2* triple mutant (*n* = 180), compared with the Col-0 control (*n* = 276). **(B)** 3D reconstructions of cells and vacuoles and quantification of vacuolar occupancy in Col-0 control (*n* = 9), *net3c* (*n* = 5) and *net3c net4a net4b #2* (*n* = 10). Scale bars in (A) and (B) = 5 µm. **(C)** Root growth of 6-days-old Arabidopsis seedlings grown on half-strength MS medium without sucrose. Col-0 control, *n* = 28; *net3c net4a net4b #2*, *n* = 35. A representative experiment is shown. Scale bar = 1 cm. **(D)** Root length of the complementation line pNET4B::NET4B-mCherry in *net3c net4a net4b* (*n* = 56) compared with Col-0 control (*n* = 58). Box limits indicate the 25th and 75th percentiles, the horizontal line indicates the median, and whiskers extend from the minimum to the maximum. Significant differences were analyzed by one-way ANOVA followed by Dunnett’s post hoc test in (A) and (B), or by Student’s *t*-test in (C) and (D). ns = not significant; \*\*\**P* < 0.001.

### NET4A alters actin organization in root cells

The observed vacuolar defects upon simultaneous loss of NET3C and NET4A/B, together with the established association of NET proteins with actin (Deeks et al., 2012; Kaiser et al., 2019), prompted us to examine whether NET4-dependent vacuolar organization is linked to the actin cytoskeleton. We therefore crossed NET4A-GFP^OE^ plants with a lifeact-RFP reporter line and investigated actin organization.

NET4A overexpression altered the organization of cortical actin filaments, with filaments showing increased orientation toward the cell center and nucleus and reduced parallel alignment (Fig. 2A and 2B). Quantification revealed significant differences in actin organization in cells of the transition zone, where actin bundles from the cell periphery were rearranged towards the nucleus (Fig. 2C). Accordingly, an impact on vacuole shape could be observed in that area. Accumulation of actin filaments in the cell center led to a shift of the vacuole towards the edges. Only thin tubular strands remained around the nucleus, keeping the two vacuolar entities in the lower and upper part of the cell connected (Fig. S4). Notably, NET4A signals seem to be more pronounced in this area and colocalize well with the actin marker lifeact-RFP (Fig. 2D). Taken together, these observations indicate that changes in NET4A abundance affect both actin organization and vacuolar morphology, consistent with the proposed role of NET proteins as membrane-associated actin regulators.

**Fig. 2.**
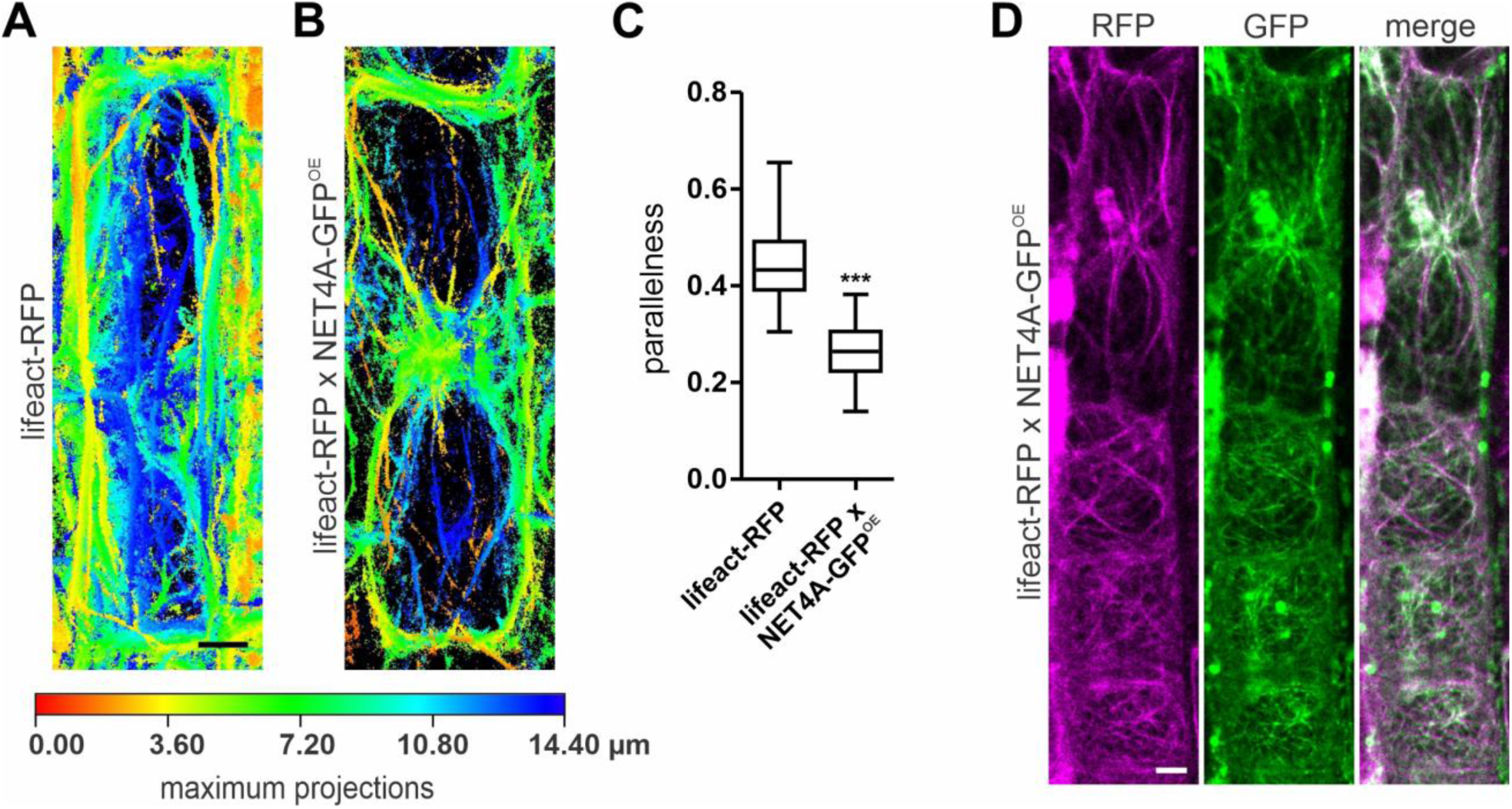
NET4 overexpression alters actin cytoskeleton organization. **(A, B)** Depth-coded maximum-intensity projections of actin filaments visualized using lifeact-RFP in representative epidermal cells from the transition zone of Arabidopsis roots under control conditions (A) and following NET4A overexpression (B). **(C)** Quantification of actin filament parallelness in plants expressing lifeact-RFP alone (*n* = 18) or together with NET4A-GFP (NET4A-GFP^OE^; *n* = 24). Statistical significance was assessed using Student’s *t*-test; \*\*\**P* < 0.001. **(D)** Colocalization of NET4A-GFP (green) and lifeact-RFP (magenta). Scale bars in all images = 5 µm. Box limits indicate the 25th and 75th percentiles, the horizontal line indicates the median, and whiskers extend from the minimum to the maximum.

### NET4A overexpression partially restores actin-dependent vacuolar defects

To further examine the relationship between NET4A and actin-dependent vacuolar organization, we introduced NET4A-GFP^OE^ into the *act7-4* mutant background. The *act7-4* mutant has previously been reported to display altered vacuolar morphology, impaired root growth, and defects in root cell organization (Gilliland et al., 2003; Scheuring et al., 2016). When NET4A was overexpressed in the *act7-4* background, irregular vacuoles of the mutant in root meristem cells partially recovered (Fig. 3A) and also the observed cell wall irregularities were fewer (Fig. 3B). Interestingly, recovery of the *act7-4* mutant phenotype by overexpression of NET4A could also be observed at the organ level, indicated by longer roots of *act7-4* x NET4A-GFP^OE^ compared to *act7-4* (Fig. 3C). These findings are consistent with a role of NET4A in coupling actin organization to vacuolar architecture.

**Fig. 3.**
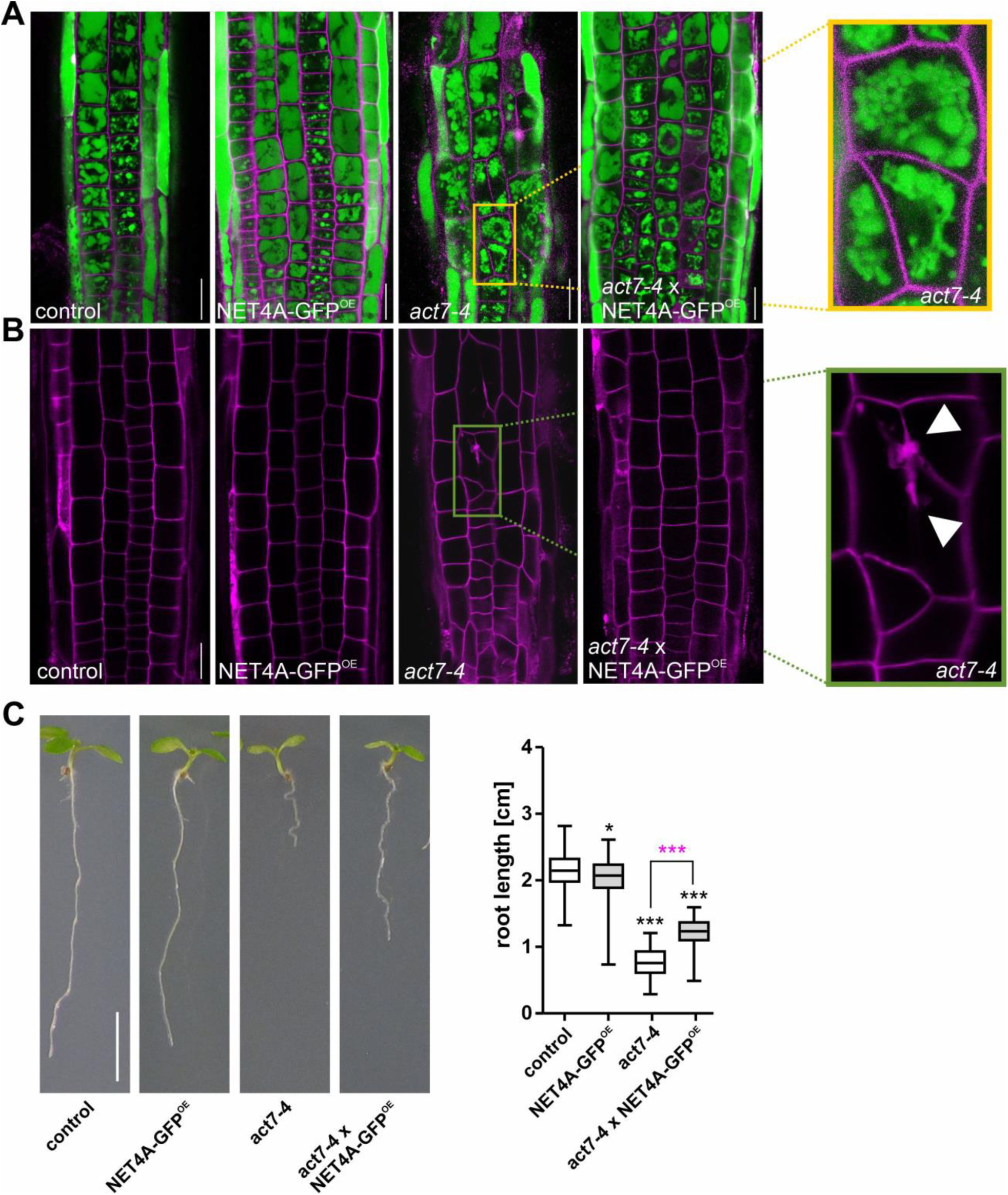
NET4A overexpression partially rescues actin-related phenotypes. **(A)** Vacuolar morphology in Arabidopsis root epidermal cells of Col-0 control, plants overexpressing NET4A-GFP (NET4A-GFP^OE^), the *act7-4* mutant, and the *act7-4* x NET4A-GFP^OE^ crossing line. Vacuoles were stained with BCECF (green), and cell walls were counterstained with propidium iodide (PI; magenta). **(B)** Cellular organization of the root epidermis from the late meristematic zone to the early elongation zone. Scale bars in (A) and (B) = 20 µm. **(C)** Root length in the Col-0 control (*n* = 153), NET4A-GFP^OE^ (*n* = 154), *act7-4* (*n* = 48), and *act7-4* x NET4A-GFP^OE^ (*n* = 59) backgrounds. Scale bar = 5 mm. Statistical significance was assessed by one-way ANOVA followed by Tukey’s post hoc test. Significant differences relative to the wild-type control are indicated; selected pairwise differences between lines are highlighted in magenta. \**P* < 0.05; \*\*\**P* < 0.001. Box limits in all graphs indicate the 25th and 75th percentiles, the horizontal line indicates the median, and whiskers extend from the minimum to the maximum.

### NET4A promotes recovery of vacuolar continuity after actin depolymerization

Since the vacuolar defects observed in *act7-4* resemble those induced by pharmacological disruption of actin filaments, we next examined whether NET4 influences vacuolar organization during actin depolymerization. Latrunculin B (LatB) treatment induces fragmentation of the vacuolar compartment into numerous small and disconnected structures (Scheuring et al., 2016). While LatB induced fragmentation of vacuoles in the control and in *net4a net4b*, NET4A-GFP^OE^ plants showed reduced vacuolar fragmentation, displaying vacuoles of almost unchanged size and shape (Fig. 4A and 4B). When monitoring lifeact-RFP during LatB treatment, we observed a seemingly slower actin filament depolymerization in the NET4A-GFP^OE^ background (Fig. S5). These observations raised the possibility that NET4A influences either the maintenance or the recovery of vacuolar organization following actin disruption. To differentiate this, we used a high LatB concentration (5 µM) for a prolonged time (16h) to entirely fragment vacuoles in Col-0 and NET4A-GFP^OE^ and performed washout experiments. Two hours after transfer to LatB-free medium, vacuolar continuity was restored in both lines, as indicated by the appearance of larger and more connected vacuolar structures (Fig. 4C). However, vacuoles in the NET4A overexpression line recovered more rapidly from fragmentation compared with wild type, consistent with a role for NET4A in vacuolar remodeling after actin disruption.

**Fig. 4.**
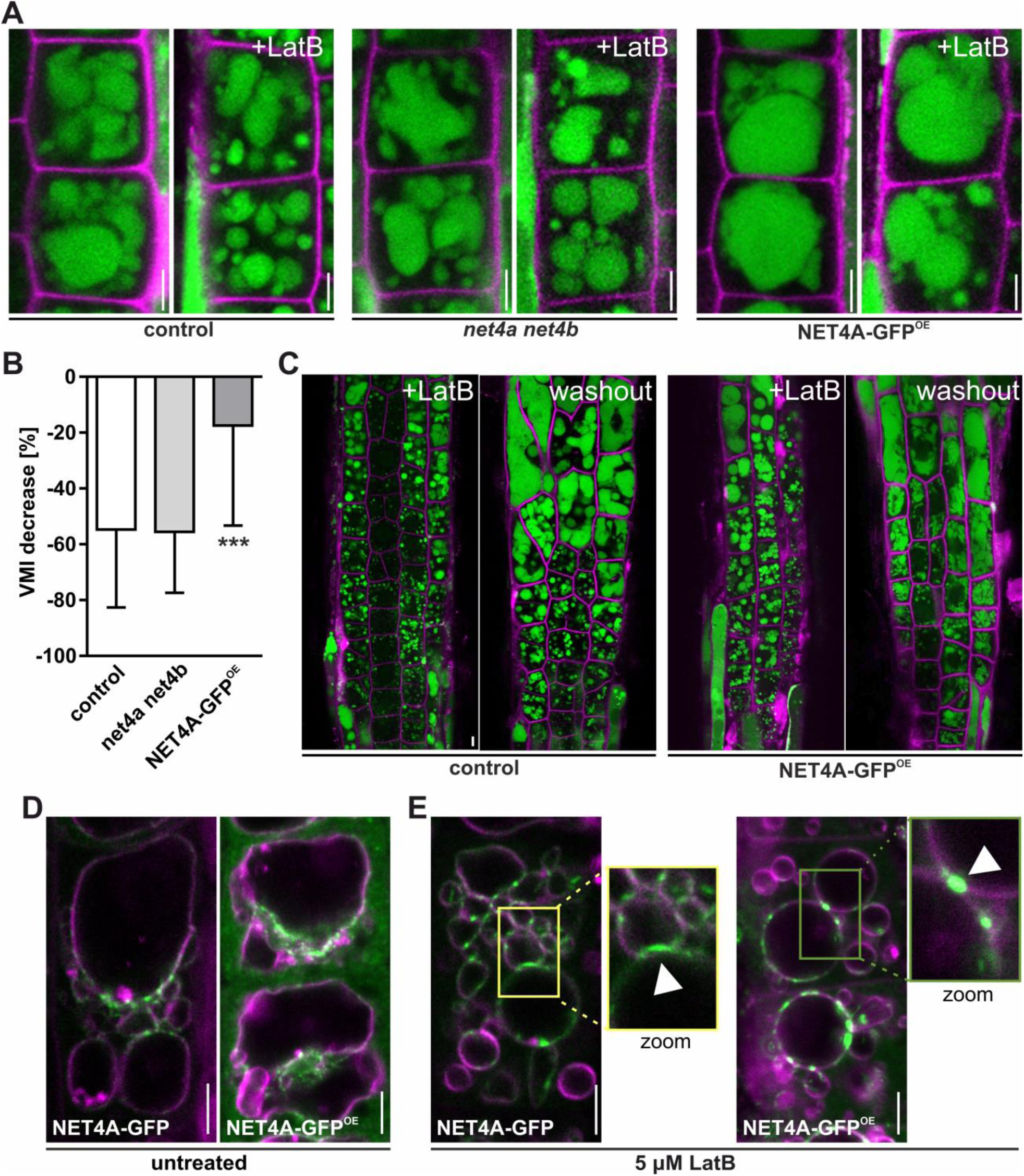
NET4A promotes vacuolar remodeling and accumulates at closely apposed vacuolar membranes. **(A)** Vacuolar morphology in Col-0 (control), the *net4a net4b* double mutant, and NET4A-GFP^OE^ plants following treatment with 500 nM latrunculin B (LatB) for 5 h. **(B)** Quantification of (A). Vacuolar morphology was quantified using the vacuolar morphology index (VMI) and changes upon LatB treatment were compared with the corresponding untreated lines as VMI decrease in percent. Col-0, *n* = 97 cells from 20 seedlings; *net4a net4b*, *n* = 86 cells from 20 seedlings; NET4A-GFP^OE^, *n* = 112 cells from 18 seedlings. **(C)** Vacuolar morphology following actin depolymerization with 5 µM LatB for 16 h and subsequent monitoring of vacuolar recovery during a 2h washout period in Col-0 and NET4A-GFP^OE^ plants. **(D)** Tonoplast staining with FM4-64 in plants expressing NET4A-GFP under its endogenous promoter or overexpressing NET4A-GFP under the CMV 35S promoter. **(E)** NET4 localization following treatment with 5 µM LatB for 5 h. Arrowheads indicate putative vacuole–vacuole fusion sites. Scale bars = 5 µm. Statistical significance relative to the Col-0 control in (B) was assessed by one-way ANOVA followed by Dunnett’s post hoc test. Bar chart columns indicate means, and error bars indicate standard deviations. \*\*\**P* < 0.001.

To investigate NET4 localization under actin-depleting conditions, we used LatB on NET4 lines under its endogenous promotor (NET4A-GFP) and under the CMV 35S promotor (NET4A-GFP^OE^). Since endogenous NET4A expression only starts in elongation cells (Kaiser et al., 2019), we investigated epidermal cells in the transition area from meristem to elongation zone. While NET4 in both lines localized in a bead-on-a-string pattern at the tonoplast and was already pronounced at apparently neighboring membrane parts (Fig. 4D), LatB treatment caused NET4 signals to accumulate at discrete tonoplast-associated foci, which frequently coincided with regions where vacuolar substructures appeared closely apposed (Fig. 4E). This suggests that NET4A is enriched at specialized tonoplast domains that may contribute to vacuolar membrane remodeling.

### NET proteins associate with RABG3/HOPS-dependent vacuolar organization

The accumulation of NET4A at closely apposed vacuolar membranes prompted us to examine its relationship with the machinery that regulates vacuolar membrane organization. In yeast and mammals, vacuolar and lysosomal membrane fusion is controlled by Rab GTPases and HOPS, which coordinate membrane tethering and SNARE-dependent fusion. In plants, RABG3 and HOPS/CORVET similarly regulate vacuolar trafficking and organization. We therefore asked whether NET4 is functionally connected to RABG3- and HOPS-dependent vacuolar organization. We first examined the genetic relationship between NET4 and the HOPS/CORVET machinery. Dexamethasone-induced expression of a VPS16 artificial microRNA (amiR-VPS16) resulted in fragmented vacuoles, similar to the phenotype previously reported following amiR-VPS39-mediated depletion of VPS39 (Takemoto et al., 2018). In contrast, vacuolar fragmentation was significantly reduced in plants expressing amiR-VPS16 in the NET4A-GFP overexpression background (Fig. 5A and 5B), indicating that increased NET4A abundance modifies the vacuolar phenotype caused by reduced VPS16 function.

**Fig. 5.**
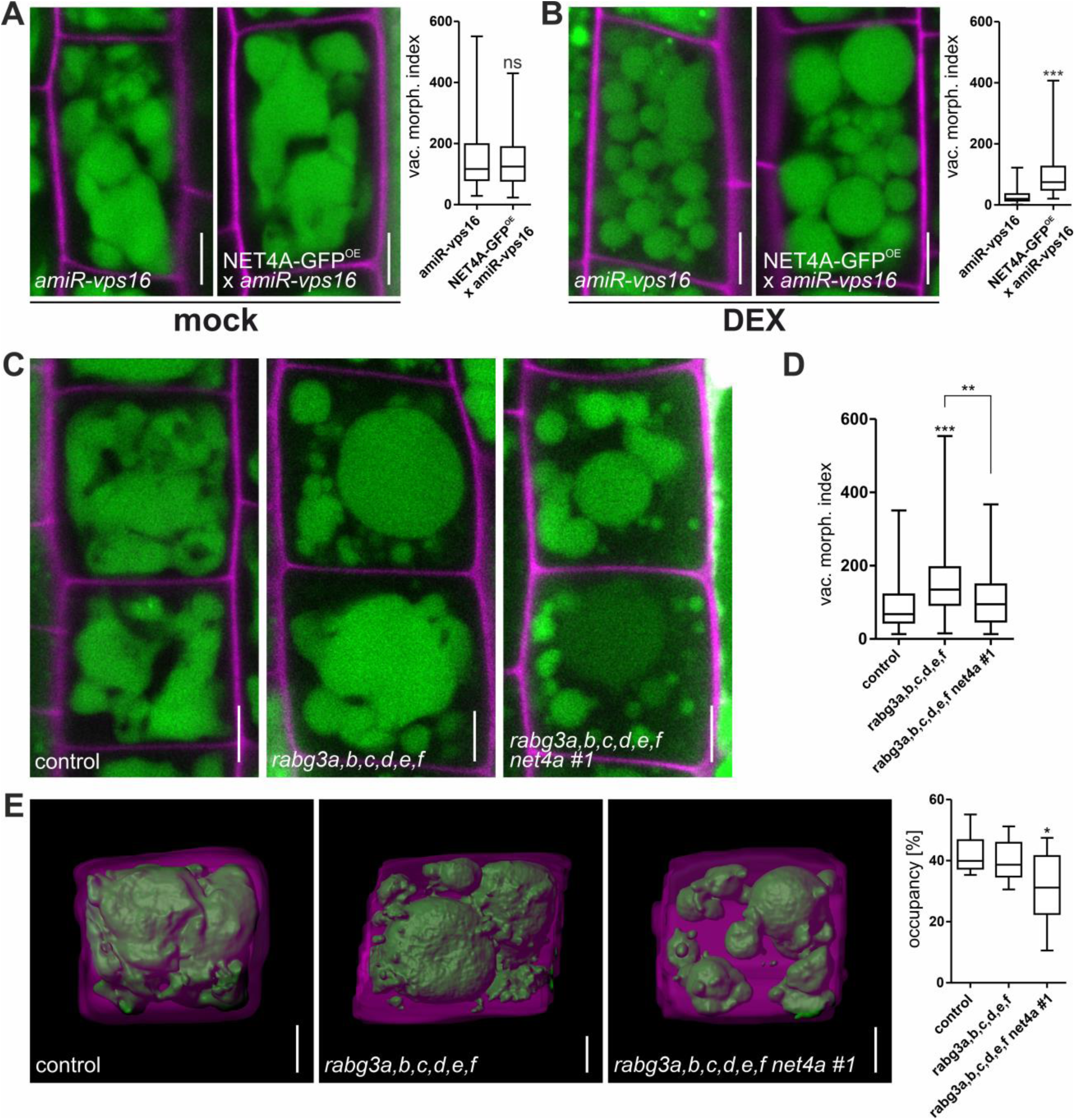
NET4 modulates vacuolar morphology in HOPS- and RABG3-deficient backgrounds. **(A, B)** Vacuolar morphology, visualized by BCECF staining, and quantification of the vacuolar morphology index (VMI) in plants expressing an artificial microRNA targeting VPS16 (amiR-vps16), a subunit shared by the HOPS and CORVET complexes, in the presence or absence of NET4A-GFP overexpression (NET4A-GFPᴼᴱ). Plants were analyzed under mock conditions (A; n = 109 and 107, respectively) or following dexamethasone (DEX) treatment (B; *n* = 100 and 97, respectively). **(C)** Vacuolar morphology in the *rabg3a,b,c,d,e,f* sextuple mutant and the *rabg3a,b,c,d,e,f net4a #1* septuple mutant compared with Col-0. **(D)** Quantification of vacuolar morphology in (C) using the VMI. Col-0, *n* = 96; *rabg3a,b,c,d,e,f*, *n* = 100; *rabg3a,b,c,d,e,f net4a #1*, *n* = 92. **(E)** 3D reconstructions of cells and vacuoles and quantification of vacuolar occupancy in Col-0 (*n* = 8), *rabg3a,b,c,d,e,f* (*n* = 9), and *rabg3a,b,c,d,e,f net4a #1* (*n* = 9). Box limits indicate the 25th and 75th percentiles, the horizontal line indicates the median, and whiskers extend from the minimum to the maximum. Statistical significance was assessed using Mann–Whitney tests in (A) and (B), and one-way ANOVA followed by Tukey’s post hoc test in (D) and (E). \**P* < 0.05; \*\**P* < 0.01; \*\*\**P* < 0.001. Scale bars = 5 µm.

Since RABG3 proteins interact with the VPS39 subunit of HOPS (Takemoto et al., 2018), we next examined the relationship between NET4 and RABG3. We analyzed the previously characterized *rabg3a,b,c,d,e,f* sextuple mutant (Ebine et al., 2014), which displayed altered vacuolar morphology, including large vacuolar bodies in the cell center accompanied by numerous smaller vacuolar structures at the cell periphery (Fig. 5C and 5D). The vacuolar morphology index was increased in the sextuple mutant, consistent with altered vacuolar organization. We then generated two independent *rabg3a,b,c,d,e,f net4a* septuple mutants by CRISPR/Cas9-mediated deletion of large portions of *NET4A* in the *rabg3* sextuple background (Fig. S1). Both septuple mutants exhibited similar but more pronounced vacuolar defects, with numerous small vacuolar structures surrounding the central vacuolar compartment. In some cells, the large central vacuole was replaced by two or more smaller vacuolar compartments (Fig. 5C and 5D; Fig. S6A). Root growth was strongly reduced in both the sextuple and septuple mutants (Fig. S6B). To further characterize the vacuolar phenotype, we generated three-dimensional reconstructions of vacuoles in root epidermal cells. Whereas control cells contained largely interconnected vacuolar structures, as reported previously (Scheuring et al., 2016), the *rabg3* sextuple mutant displayed numerous disconnected vacuolar substructures. These defects were further enhanced in the *rabg3 net4a* septuple mutants, in which the disconnected vacuolar structures occupied a smaller fraction of the cell (Fig. 5E). Together, these genetic interactions indicate that NET4 contributes to the organization of the vacuolar compartment in the context of RABG3- and HOPS-dependent trafficking.

### NET proteins associate with RABG3 GTPases

To investigate the molecular basis of the genetic relationship between NET4 and RABG3/HOPS-dependent vacuolar organization, we performed co-immunoprecipitation followed by LC-MS/MS analysis using NET4A-GFP expressed from its endogenous promoter and NET4A-GFP overexpressed from the 35S promoter. Numerous proteins were enriched in NET4A-GFP immunoprecipitates relative to the corresponding wild type controls (Fig. S7A and S7B; Table S1). Among the significantly enriched proteins were RABC1 and two RABG3 GTPases, together with several additional proteins. The identification of RABG3 proteins in NET4A-GFP immunoprecipitates was consistent with previous evidence for an association between NET4 and RABG3 GTPases (Hawkins et al., 2023). We could confirm these interactions by using rBiFC and mating-based split-ubiquitin system (mbSUS) (Grefen et al., 2009) assays (Fig. S7). Interestingly, RABG3 association was not restricted to NET4: we also detected interaction of RABG3 proteins with NET3A and NET3C (Fig. 6A; Fig. S8). This finding extends the previously described NET4–RABG3 association to NET3 proteins and is consistent with the presence of the conserved IRQ domain in NET3A and NET3C, which has been implicated in RABG3 binding in NET4 proteins (Hawkins et al., 2023).

**Fig. 6.**
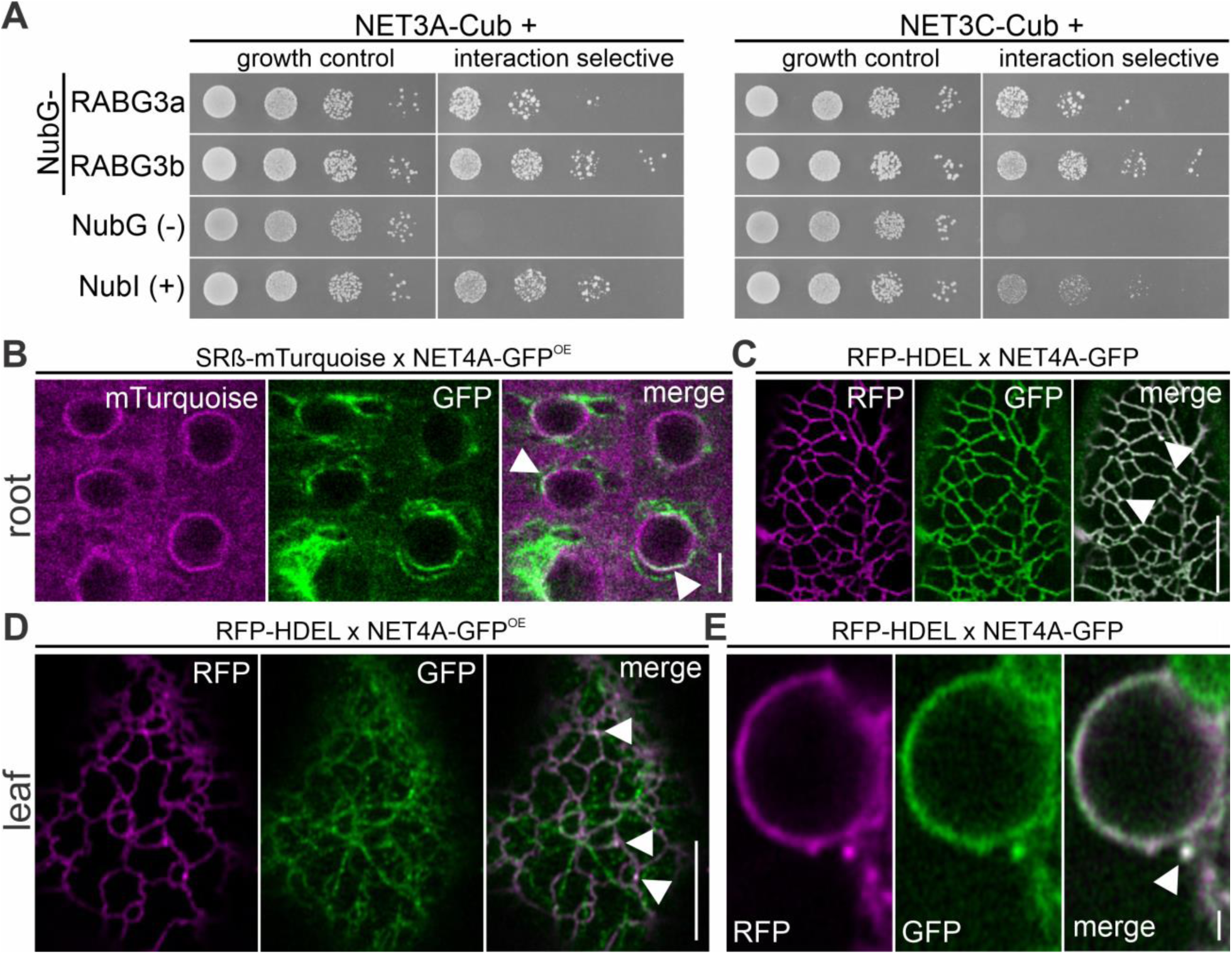
NET3 proteins associate with RABG3 GTPases, and NET4A localizes to the endoplasmic reticulum. **(A)** Mating-based split-ubiquitin system (mbSUS) growth assays testing the association of NET3A-Cub and NET3C-Cub with NubG-RABG3a and NubG-RABG3b in yeast. THY.AP4 (MATa) strains expressing Cub fusions were mated with THY.AP5 (MATα) strains expressing NubG fusions. Serial dilutions (OD₆₀₀ = 1 to 0.001) were spotted onto control medium and interaction-selective medium. Growth on control medium was assessed after 36 h at 28°C to confirm the presence of the fusion constructs; growth on selective medium was assessed after 72 h to test for interaction-dependent reporter activation. NubG and NubI served as negative and positive controls, respectively. **(B)** Colocalization of overexpressed NET4A-GFP (NET4A-GFP^OE^) with the ER marker SRβ-mTurquoise in Arabidopsis roots. **(C)** Colocalization of NET4A-GFP expressed under its endogenous promoter with the ER marker RFP-HDEL in Arabidopsis roots. **(D)** Colocalization of NET4A-GFP^OE^ with RFP-HDEL in Arabidopsis leaves. **(E)** Colocalization of NET4A-GFP expressed under its endogenous promoter with RFP-HDEL in Arabidopsis leaves. Arrowheads indicate colocalization. Scale bars = 5 µm in (B–D) and 1 µm in (E).

### NET proteins associate with ER-associated membrane domains

NET3 proteins have previously been associated with the ER and implicated in cytoskeleton-associated organization of ER–plasma membrane interfaces (Wang et al., 2014; Zang et al., 2021). Given the overlapping functions of NET3 and NET4 in vacuolar organization, we asked whether NET4 proteins also associate with the ER. To determine this, we first co-infiltrated NET4A-mTurquoise and NET4B-GFP expression constructs together with an RFP-HDEL ER marker construct into tobacco leaf epidermal cells. Here, we found partial co-localization of both NET4 proteins with the ER marker, especially at the nuclear envelope (Fig. S9). We further crossed Arabidopsis NET4A-GFP and NET4A-GFP^OE^ lines with the ER markers RFP-HDEL or SRß-mTurquoise (the β-subunit of the ER membrane-associated signal recognition particle receptor). Using high- and super-resolution microscopy, we could confirm NET4A localization at the nuclear envelope and at the ER in roots (Fig. 6B and 6C) and in leaves (Fig. 6D and 6E). Notably, NET4A signals at the ER were often found enhanced in punctate signals (indicated by arrowheads), resembling the vacuole connection sites we observed earlier (Fig. 4D). The somewhat surprising ER-associated localization of NET4A raised the possibility that NET4 proteins are components of ER membrane domains involved in interorganelle communication.

Since it has been shown that NET3C recruitment to the ER is mediated by its interaction with the ER-integral membrane contact site marker VAP27-1 (Wang et al., 2014), we tested whether NET4s can also interact with VAP27-1. Yeast mbSUS assays confirmed the interaction of NET4A and NET4B with VAP27-1, supporting evidence for their presence at the ER (Fig. 7A; Fig. S8). To screen for meeting points of VAP27-1 and NET4A *in vivo*, we used an approach based on the split super-folder GFP system (Cabantous et al., 2005). Thus, we generated constructs harboring two expression cassettes. In one of these, the sequence coding for the first 10 β-strands of GFP is fused to VAP27-1 (GFP1-10-VAP27-1) and expression controlled by a constitutive promoter. In the other cassette, NET4A is tagged with the 11^th^ GFP β-strand (NET4A-GFP11). To avoid constitutive expression of both fusions, we used an estradiol-inducible promoter (Denninger et al., 2019). Since both split-GFP parts alone are non-fluorescent, only at sites where VAP27-1 and NET4A meet, the two GFP parts should be reconstituted, emitting fluorescence upon excitation (Fig. 7B). Using transient expression in tobacco leaves, we expressed GFP1-10-VAP27-1 and NET4A-GFP11 together with the ER marker RFP-HDEL (Fig. 7B). Four days after induction of estradiol-induced expression, we observed punctate GFP-signals at the ER, representing potential VAP27-1 and NET4A contact sites.

**Fig. 7.**
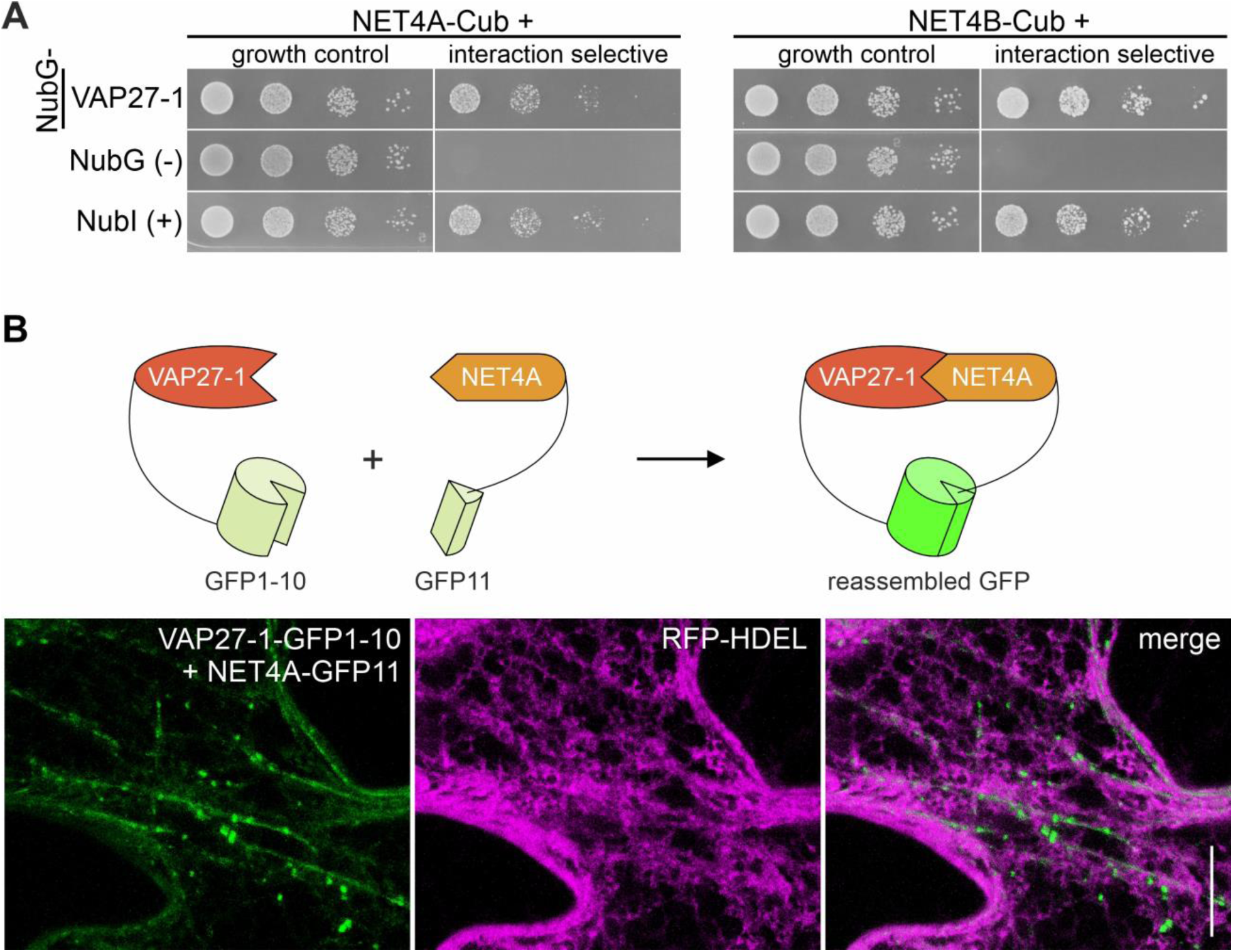
NET4 associates with the ER membrane contact site protein VAP27-1. **(A)** Mating-based split-ubiquitin system (mbSUS) growth assays testing the association of NET4A-Cub and NET4B-Cub with NubG-VAP27-1 in yeast. THY.AP4 (MATa) strains expressing Cub fusions were mated with THY.AP5 (MATα) strains expressing NubG fusions. Serial dilutions (OD₆₀₀ = 1 to 0.001) were spotted onto control medium and interaction-selective medium. Growth on control medium was assessed after 36 h at 28°C to confirm the presence of the fusion constructs; growth on selective medium was assessed after 72 h to test for interaction-dependent reporter activation. NubG and NubI served as negative and positive controls, respectively. **(B)** Principle of the split-GFP assay and colocalization of GFP1–10-VAP27-1and NET4A-GFP11 with the ER marker RFP-HDEL following transient expression in *Nicotiana benthamiana*. The two GFP fragments are non-fluorescent when expressed separately; when brought into proximity, they reconstitute a fluorescent GFP signal. A 3D projection of z-stack images is shown.

### NET4B interacts with ORP2A and NET proteins contribute to membrane lipid homeostasis

Since NET4 interacts with VAP27-1 at the ER (Fig. 7), we asked whether this association might connect NET proteins to ER–vacuole lipid homeostasis. Membrane contact sites have been implicated in the regulation of vacuolar morphology through lipid metabolism, including sphingolipid metabolism at the nucleus–vacuole junction in yeast (Hanaoka et al., 2024). We therefore investigated whether NET4 proteins associate with lipid transfer proteins that could link ER membrane domains to the vacuole. Using the mating-based split-ubiquitin system (mbSUS), we found that NET4B associated with the ER-localized OSBP-related protein ORP2A, whereas no association was detected with the other tested lipid transfer proteins (Fig. 8A; Fig. S10A).

**Fig. 8.**
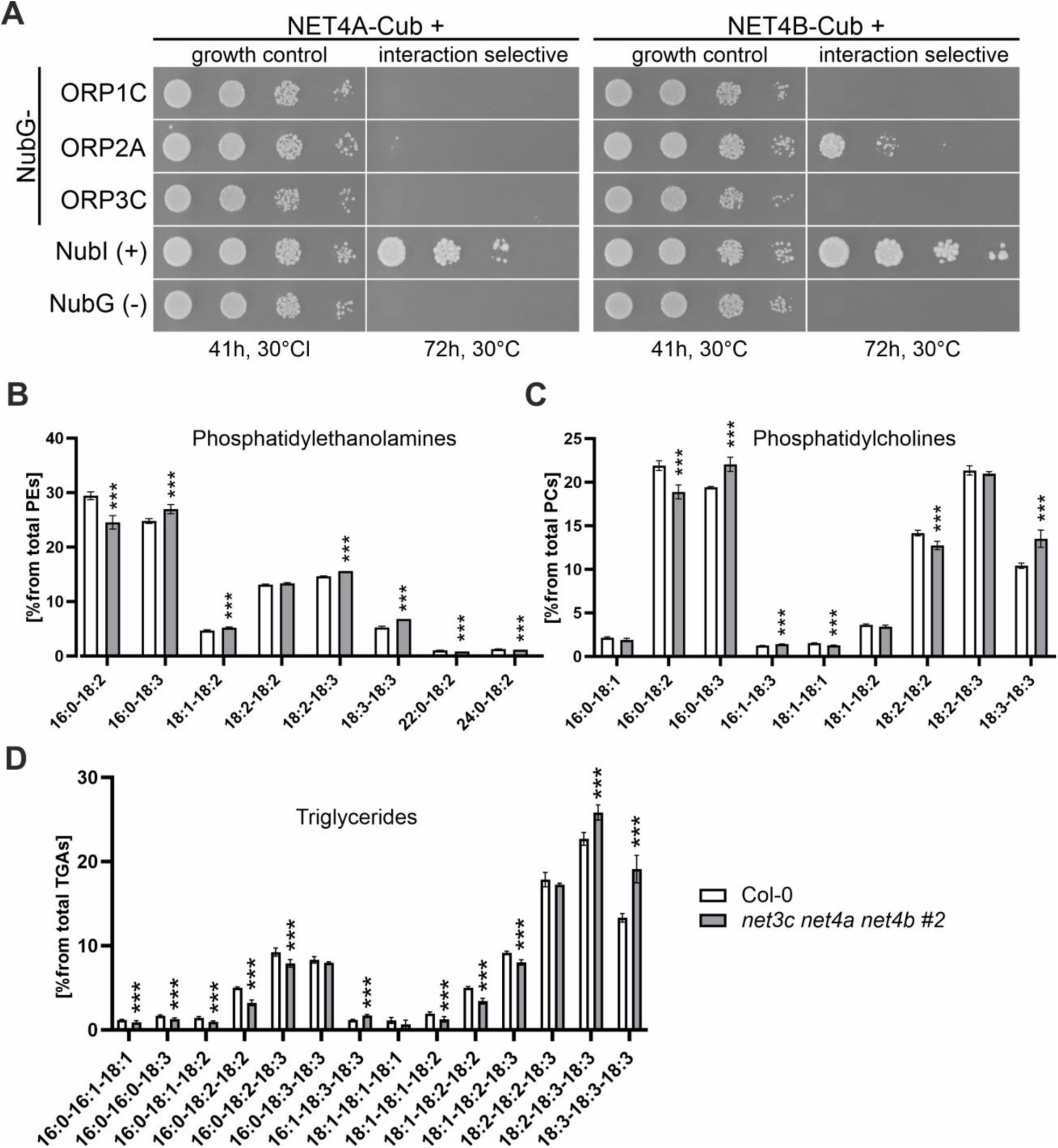
NET3 and NET4 proteins contribute to lipid homeostasis. **(A)** Mating-based split-ubiquitin system (mbSUS) growth assays testing the association of NET4A-Cub and NET4B-Cub with NubG-ORP1C, NubG-ORP2A, and NubG-ORP3C in yeast. THY.AP4 (MATa) strains expressing Cub fusions were mated with THY.AP5 (MATα) strains expressing NubG fusions. Serial dilutions (OD₆₀₀ = 1 to 0.001) were spotted onto control medium and interaction-selective medium to confirm the presence of fusion constructs and test for interaction-dependent reporter activation, respectively. The incubation times and temperature are as indicated in the figure. NubG and NubI served as negative and positive controls, respectively. **(B–D)** LC–MS/MS-based lipidomic analysis comparing Col-0 and the *net3c net4a net4b* triple mutant (*n* = 4 per genotype). Only lipid species accounting for more than 1% of the total species within their respective lipid classes are shown. See Supplementary Dataset 1 for the complete list of detected lipid species. Statistical differences were assessed using Student’s *t*-tests. \**P* < 0.05; \*\**P* < 0.01; \*\*\**P* < 0.001.

The association of NET4B with ORP2A prompted us to examine whether loss of NET proteins affects membrane lipid composition. We performed lipidomic analyses of the *net3c net4a net4b* triple mutant and Col-0. The triple mutant displayed significant changes in several lipid species, including pronounced alterations in phosphatidylethanolamine (PE) and phosphatidylcholine (PC) species as well as triglycerides (Fig. 8B-8D). These changes indicate altered membrane lipid homeostasis in the absence of NET3 and NET4 proteins. To further examine the relationship between NET proteins and sphingolipid-dependent processes, we treated Col-0 and *net3c net4a net4b* seedlings with 1 µM fumonisin B1, an inhibitor of sphingolipid biosynthesis (Zhao et al., 2022). Fumonisin B1 reduced root growth in both genotypes; however, the *net3c net4a net4b* mutant was significantly less affected than Col-0 (Fig. S10B-D). Together, these findings associate NET4B with the lipid transfer protein ORP2A and indicate that NET proteins contribute to the maintenance of membrane lipid homeostasis. Integrating our findings, we propose that NET proteins act as adaptable membrane-associated organizers that connect the actin cytoskeleton with distinct membrane processes (Fig. 9).

**Fig. 9.**
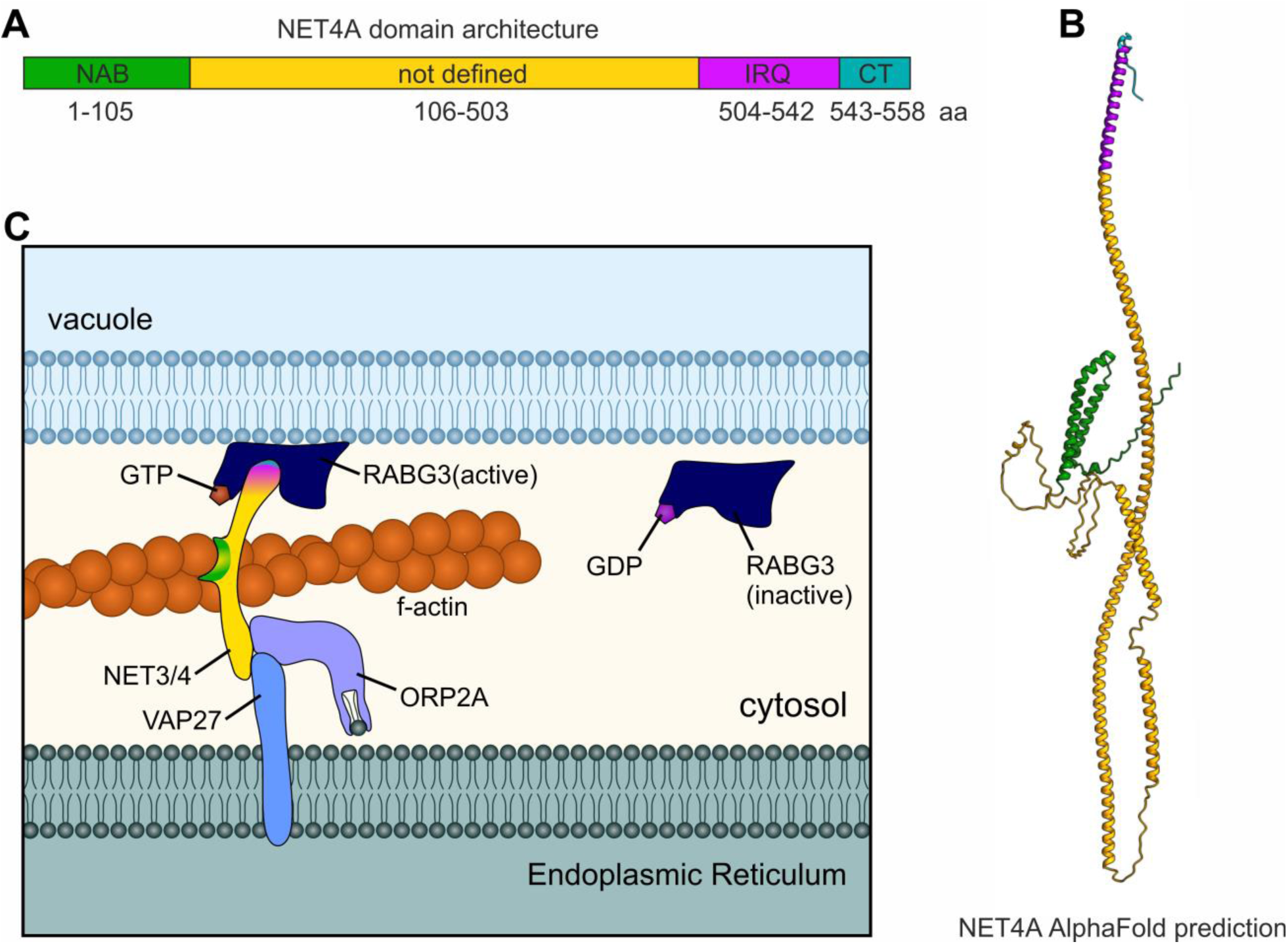
NET4A domain architecture and proposed model of NET protein functions at the ER–vacuole interface. **(A)** Schematic representation of the NET4A domain architecture, showing the NAB domain (amino acids 1–105), a region of undefined domain architecture (amino acids 106–503), the IRQ domain (amino acids 504–542), and the C-terminal region (CT; amino acids 543–558). **(B)** AlphaFold-predicted structure of NET4A, with regions colored according to the domain architecture shown in (A). **(C)** Proposed model illustrating how NET3/NET4 proteins may link F-actin organization to vacuolar membrane processes and ER-associated membrane contact sites. NET3/NET4 proteins associate with active RABG3 GTPases at the vacuolar membrane and with the ER membrane contact site protein VAP27-1. The association of NET4B with the lipid-transfer protein ORP2A further links NET proteins to membrane lipid homeostasis. These interactions may contribute to the coordination of actin organization, vacuolar remodeling, and membrane-associated processes at the ER–vacuole interface. Active RABG3 is shown in its GTP-bound state and inactive RABG3 in its GDP-bound state. The model illustrates potential functional relationships and does not imply that all depicted proteins necessarily form a single molecular complex.

## Discussion

In this study, we aimed to elucidate the molecular mechanisms underlying actin-dependent remodeling of the plant vacuole. By integrating cell biological, molecular, and biochemical approaches, we identified members of the actin-binding NET (NETWORKED) protein family as key factors linking actin organization to vacuolar architecture. Our data indicates that NET3 and NET4 proteins have overlapping functions in the organization of the vacuole and that NET4 connects actin organization with vacuolar remodeling. In addition, our genetic and interaction analyses associate NET3 and NET4 proteins with RABG3 GTPases, ER-associated membrane domains, and lipid homeostasis. Together, these findings suggest that NET proteins act as membrane-associated adaptors that coordinate the actin cytoskeleton with distinct membrane processes.

### NET3 and NET4 redundantly regulate vacuolar architecture

Our previous findings indicate that the vacuole-cytoskeleton interface is crucial for regulating vacuolar morphology and size (Scheuring et al., 2016). Here, the analysis of higher-order NET mutants demonstrates that NET3 and NET4 proteins have overlapping functions in this process. While the *net4a net4b* double mutant has only a limited phenotype (Kaiser et al., 2019; Hawkins et al., 2023), combining *net3* and *net4* mutations resulted in pronounced defects in vacuolar organization and plant growth. The presence of several smaller vacuolar structures that appeared to disconnect from the main vacuolar body, together with reduced vacuolar occupancy in the *net3c net4a net4b* triple mutant, indicates a strong disturbance of vacuolar organization.

The functional redundancy between *NET3* and *NET4* might explain the lack of a pronounced and global *net4a net4b* phenotype. While none of the other NET superfamily members were transcriptionally upregulated in the *net4a net4b* background (Kaiser et al., 2021), compensation could be achieved by NET3 proteins at the protein level. Conversely, the similar effects of NET4A and NET4B overexpression on vacuolar occupancy and root growth are consistent with overlapping functions among NET family members. Thus, the combined genetic and overexpression data support a model in which different NET proteins contribute to a common membrane–actin regulatory system, with individual members providing partially redundant functions. The reduced vacuolar occupancy in the higher-order mutant is likely to contribute to the impaired root growth, since restricting vacuolar size interferes with the space-filling function of the vacuole that is required for rapid cell elongation (Kaiser and Scheuring, 2020). These findings therefore connect NET-dependent organization of the vacuole to plant growth and provide a framework for understanding how the actin cytoskeleton contributes to vacuolar expansion.

### NET4 connects actin organization with vacuolar remodeling

Our data indicates that NET4 abundance affects both, actin organization and vacuolar architecture. More compact vacuoles in NET4A overexpression plants were accompanied by actin filaments redirected towards the nucleus. Thus, in addition to the original description that NET4 accumulates at sites of vacuolar constrictions (Kaiser et al., 2019), NET4 appears to be particularly enriched at discrete regions of the tonoplast where actin organization and vacuolar architecture are closely associated. Overexpression of NET4A partially compensated for the loss of vegetative actin *ACT7*, supporting a functional connection between NET4-dependent membrane association and actin organization (Fig. 3). ACT7 has been shown to be specifically involved in root growth and architecture and to be highly responsive to treatment with the phytohormone auxin (McDowell et al., 1996; Kandasamy et al., 2009). Although actin filaments recruited towards the nuclear envelope by NET4A-GFP were in close proximity to the nucleus, we could not detect changes in nuclear positioning in NET4A overexpression or *net4a net4b* plants. Thus, the altered actin organization is more likely to reflect a role of NET4 in membrane-associated actin organization than a specific function in nuclear positioning. Limited vacuole fragmentation and faster recovery of vacuolar continuity after LatB washout further suggest that NET4 contributes to actin-dependent vacuolar remodeling. Upon actin depolymerization, NET4 signals remained associated with discrete regions of the tonoplast, including sites where vacuolar structures remained closely apposed. These observations are reminiscent of the specialized membrane regions observed during vacuole fusion in yeast, where actin and membrane-associated factors accumulate at sites of vacuolar docking and fusion (Eitzen et al., 2002; Starr and Fratti, 2019). In yeast, vacuole fusion involves the coordinated action of Rab GTPases, HOPS, SNARE proteins, and the actin cytoskeleton (Wickner, 2010; Balderhaar and Ungermann, 2013). Our data does not establish NET4 as a component of the fusion machinery itself, but suggests that NET4 may organize actin at membrane domains that are important for vacuolar remodeling.

### NET proteins associate with RABG3-dependent vacuolar organization

The association of NET proteins with RABG3 GTPases provides a molecular connection between NET-dependent actin organization and the machinery controlling vacuolar membrane identity and remodeling. NET4A and NET4B have previously been shown to interact with active RABG3 GTPases through their IRQ domain (Hawkins et al., 2023). Our findings extend this by demonstrating that NET3A and NET3C, also possessing an IRQ domain, associated with RABG3 proteins as well. Thus, RABG3 interaction is not restricted to the NET4 proteins residing at the tonoplast but represents a broader property of NET proteins containing the conserved IRQ domain. The genetic interaction between NET4 and the RABG3/HOPS-dependent vacuolar pathway further supports a functional relationship between these proteins. NET4A overexpression partially alleviated the vacuolar phenotype caused by HOPS impairment, whereas deletion of NET4 enhanced the vacuolar defects of the *rabg3a,b,c,d,e,f* mutant background (Fig. 5). Together with the localization of NET4 at discrete tonoplast domains, these results are consistent with a role for NET proteins in organizing membrane-associated actin at sites where RABG3-dependent vacuolar trafficking or remodeling occurs. Whether NET proteins act upstream, downstream, or in parallel to RABG3/HOPS remains to be determined. This also provides a mechanistic context for the actin-dependent recovery of vacuolar continuity observed after LatB treatment. The association of NET4 with RABG3-positive membrane domains raises two, not mutually exclusive, possibilities. NET4 could function as a spatial link between the actin cytoskeleton and RABG3-dependent membrane domains, thereby facilitating the reorganization of vacuolar membranes following cytoskeletal perturbation. Alternatively, NET4 could be a component of the membrane fusion machinery itself, acting together with RABG3/HOPS-dependent factors at sites of vacuolar fusion. The localization of NET4 at closely apposed vacuolar membranes, its genetic interaction with the RABG3/HOPS pathway, and the faster recovery of vacuolar continuity after actin depolymerization allow for both models. Distinguishing whether NET4 primarily organizes actin at fusion sites or directly contributes to the fusion process itself will require future biochemical and functional analysis.

### NET proteins associate with ER-associated membrane domains

The identification of NET4 at ER-associated membrane domains suggests that the function of NET proteins extends beyond the vacuolar membrane. NET3 proteins have previously been shown to associate with the ER and nuclear envelope and to participate in cytoskeleton-dependent membrane organization (Wang et al., 2014; Zang et al., 2021). In addition, NET3C, VAP27, KLCR1, and IQD2 have been proposed to form adaptor complexes linking the cytoskeleton with ER-associated membrane domains (Zang et al., 2021). The presence of an IRQ domain in NET3A, NET3C, NET4A, and NET4B further suggests that related molecular mechanisms may operate at different membranes (Hawkins et al., 2023). Our observation that NET4 proteins associate with the ER by interacting with the ER membrane protein VAP27-1 provides additional evidence for overlapping functions of NET3 and NET4 at different membrane systems. These findings raise the possibility that NET proteins can participate in membrane-associated adaptor complexes of different composition, depending on their interaction partners and subcellular localization. In this view, the ability of NET proteins to bind actin is combined with membrane-specific interactions to organize the cytoskeleton at distinct membrane domains. The unexpected localization of NET4 proteins at the ER and their association with the MCS protein VAP27-1 support overlapping NET3–NET4 functions at different membrane systems. In yeast, the nucleus–vacuole junction (NVJ) forms a classical ER–vacuole membrane contact site through Nvj1 and Vac8 (Pan et al., 2000). While plants lack direct orthologs for this complex, functional equivalents of plant ER–vacuole MCSs are only now beginning to emerge. As illustrated in the proposed model (Fig. 9), NET4-containing complexes could contribute to the organization of ER–vacuole membrane contacts. NET proteins may link actin filaments to ER- and vacuole-associated membrane domains: their association with VAP27-1 provides a potential connection to the ER, while their association with RABG3 GTPases links them to vacuolar membrane organization. NET proteins could therefore help organize actin at closely apposed membranes or participate in larger membrane-associated complexes that stabilize or regulate these interfaces. The association of NET4B with the lipid-transfer protein ORP2A further connects this model to membrane lipid homeostasis. However, these interactions do not establish that NET4, VAP27-1, and ORP2A form a single complex or that NET proteins directly tether the ER to the vacuole. Nor do they establish whether NET4 contributes to the formation of membrane contact sites or acts at pre-existing interfaces. Instead, the model proposes that NET proteins may link actin organization with membrane-associated processes at distinct organelles, providing a framework for testing how cytoskeletal organization and membrane interactions are coordinated in plant cells.

### NET4B–ORP2A association links NET proteins to lipid homeostasis

The association of NET4B with the lipid transfer protein ORP2A provides a link between NET proteins and membrane lipid homeostasis. ORP2A and VAP27 have recently been shown to participate in the formation of an ER–chloroplast contact site, and loss of the VAP27–ORP2A complex altered lipid levels in chloroplasts (Renna et al., 2024). In our study, the *net3c net4a net4b* triple mutant displayed altered membrane lipid composition and an altered response to Fumonisin B1. These observations indicate that NET proteins contribute to the maintenance of membrane lipid homeostasis. The connection between NET proteins, VAP27-1, and ORP2A is particularly interesting in the context of vacuolar expansion. The ER can contribute membrane material during vacuole biogenesis (Viotti et al., 2013; Scheuring et al., 2024), and lipid exchange between these organelles is therefore likely to contribute to membrane growth. ER–lysosome contact sites have been described in animals and contribute to lipid exchange and organelle homeostasis (Tan and Finkel, 2022). However, whether a comparable mechanism operates between the ER and vacuole in plants remains unknown. Our data raise the possibility that NET-containing membrane-associated complexes contribute indirectly to lipid exchange between the ER and vacuole at plant MCSs. However, the altered lipid composition of the higher-order mutant and the association of NET4B with ORP2A do not by themselves demonstrate lipid transfer through NET-dependent membrane contact sites. Rather, they provide evidence that NET proteins are connected to pathways controlling membrane lipid homeostasis and establish a basis for testing whether NET-dependent membrane organization influences lipid exchange between organelles.

### NET proteins as membrane–actin organizers

Taken together, our results suggest that NET proteins function as adaptable membrane-associated organizers that connect the actin cytoskeleton with distinct membrane processes (Fig. 9C). NET3 and NET4 proteins have overlapping functions in vacuolar organization, while NET4A connects changes in actin organization with vacuolar remodeling. Their association with RABG3 GTPases links NET proteins to the machinery controlling vacuolar membrane organization, whereas their association with ER-associated proteins and ORP2A connects them to ER-associated membrane domains and lipid homeostasis. We propose that, depending on their interaction partners and subcellular localization, NET proteins form membrane-associated complexes of different compositions that couple actin filaments to distinct organelles or membrane domains. At the vacuole, NET3 and NET4 proteins may organize actin at RABG3-associated membrane domains involved in vacuolar remodeling, while potentially also contributing to the fusion process itself. At ER-associated domains, NET3/NET4-containing complexes may similarly organize actin and participate in the organization of ER–vacuole contacts. Their association with ORP2A, together with the altered lipid composition of higher-order NET mutants, further connects this model to membrane lipid homeostasis, although a direct role for NET proteins in lipid transfer remains to be established. In this model, NET proteins do not represent a single dedicated component of one membrane-remodeling pathway. Rather, they provide a potentially modular link between actin and membranes that can be adapted to different cellular contexts. The functional overlap between NET3 and NET4 may help maintain these membrane–actin connections when individual NET proteins are absent. Such coordination could be particularly important in plant cells, where vacuolar expansion is tightly coupled to cell elongation.

## Material and Methods

### Plant Material and Growth Conditions

*Arabidopsis thaliana* ecotype Columbia-0 (Col-0) was used as wild type control. The following lines were described previously: lifeact-RFP (Rosero et al., 2016), 35S::NET4A-GFP (NET4A-GFP^OE^) (Kaiser et al., 2019), NET4A::NET4A-GFP (NET4A-GFP) (Deeks et al., 2012), RFP-HDEL, *act7-4* (Gilliland et al., 2003), *net4a net4b* (Kaiser et al., 2019) and *rabg3a,b,c,d,e,f* (Ebine et al., 2014). Seeds were surface sterilized with ethanol and plates were stratified at 4 °C for 1–2 days in the dark and grown vertically at 22 °C under a 16 h light/8 h dark-cycle. For seedling growth, half-strength Murashige and Skoog (MS) medium (Duchefa, Netherlands), including 1% (w/v) sucrose (Roth, Germany), 2.56 mM MES (Duchefa) and 1% (w/v) Phytoagar (Duchefa) was used at pH 5.7.

### Chemicals and Treatments

The dyes MDY-64, propidium iodide (PI), FM4-64 and BCECF-AM (BCECF) were acquired from Life Technologies (CA, USA). The synthetic auxin α-Naphthaleneacetic acid (NAA) and dexamethasone (DEX) were obtained from Duchefa, latrunculin B (LatB) and β-estradiol from Sigma-Aldrich (MO, USA). Fumonisin B1 was obtained from abcr (Karlsruhe, Germany). PI was dissolved in water, β-estradiol in ethanol, Fumonisin B1 in methanol and all other chemicals were dissolved in dimethyl sulfoxide (DMSO).

### RNA Extraction and Quantitative Real Time PCR

RNA extraction and transcription into cDNA was carried out as described previously (Jeblick et al., 2023). For qRT-PCR, the PerfeCTa SYBR Green SuperMix for iQ (Quanta BioSciences, USA) was used and reactions were carried out in a CFX Connect Real-Time PCR Detection System (Bio-Rad, USA). Reaction procedures were set to an initial heating step of 95 °C for 3 min as well as 40 cycles of denaturing at 95 °C for 10 s followed by combined annealing-extension at 60 °C for 40 s. Primers used for target and housekeeping genes (*PP2A*) are shown in Table S2. To calculate relative expression ratios, the efficiency corrected calculation model was used (Pfaffl, 2001).

### Cloning

Expression constructs were generated using the GreenGate system (Lampropoulos et al., 2013) and are listed in Table S3. Primers used to amplify sequences for the creation of new GreenGate entry vectors are given in Table S4. For the amplification of split-GFP sequences, plasmid GFP1-10-miniCMV-GFP11×11-GFP-tDeg (Addgene plasmid # 185404 from Jian-Hui Jiang) was used (Zhou et al., 2021). Modules for GFP-, mTurquoise-, and mCherry-tags, seed coat markers and estradiol-inducible promoter (pGGD011, pGGD-GSL-mTurquoise, pGGD010, pGGF Alli-mCherry,pGGF Alli-YFP, pGGA-Ubi-XVE_oLexA-35S) were published before (Lupanga et al., 2020; Stührwohldt et al., 2020; Waadt et al., 2017; Stephani et al., 2020; Denninger et al., 2019). Internal *Bsa*I sites were mutated. To this end, additional internal primers were used to introduce suitable silent point mutations during the amplification of separate sequence parts, and the resulting PCR products were fused. In case *Bsa*I sites were present in promoter sequences, no entry vector was generated. Instead, the sequence was amplified using suitable primers to produce matching overhangs by using alternative restriction enzymes. Subsequently, the final construct was generated preassembling all modules without the promoter module. Later, the promoter sequence with matching overhangs was added by ligation in a separate reaction. A list of all entry modules used or generated in this study is provided as Table S5. The DEX-inducible amiRNA based knockdown constructs for *VPS16* (*amiR-vps16*) was cloned as described for other *VPS* (Takemoto et al., 2018). Primers used to amplify intermediate and final amiR sequences against *VPS16* are listed in Table S6. Deletion mutants were obtained using the egg cell-specific promoter-controlled CRISPR/Cas9 cloning system based on the vector pHEE401E (Wang et al., 2015). To improve the screening for transformed plants, the hygromycin resistance cassette of the original vector was replaced by a mCherry seed coat marker cassette from the GreenGate vector pGGF Alli-mCherry, resulting in the generation of pHEE-mCherry. To this end, available *EcoRI* and *SacII* restriction sites of pHEE401E were used to cut out the hygromycin resistance cassette. Vector backbone sequences that were thereby deleted in addition to the cassette were amplified by PCR and cloned back together with the PCR-amplified seed coat marker cassette. Used Primers are given in Table S7. The web tools CHOPCHOP (https://chopchop.cbu.uib.no/ (Labun et al., 2016)) and CCTop (https://cctop.cos.uni-heidelberg.de/ (Stemmer et al., 2015)) were used to select suitable guide RNAs (gRNAs). All constructs were generated by inserting a PCR fragment with two gRNAs for the same gene in the used vector as described in Xing et al. (Xing et al., 2014), aiming to delete the region between the two target sites (Fig. S1). Amplification of the respective insertion fragments was performed using the primers listed in Table S8 and vector pHEE2E-TRI as a template. All generated CRISPR/Cas9 constructs are listed in Table S9 and details about the used base vector and gRNAs are given. The obtained mutant lines are shown in Table S10.

DNA constructs for rBiFC and mbSUS assays were generated via the 2in1 cloning system (Mehlhorn et al., 2018) or Gateway^TM^ technology (Invitrogen, Thermo Fisher Scientific, Carlsbad, CA, USA). Coding sequences were PCR amplified from Col-0 seedling’s cDNA or suitable vectors harbouring the sequence. In case of At4g01245, the non-spliced gene was amplified from genomic DNA of Col-0 leaves. All primers are given in Table S11. PCR products were recombined into pDONR221-P1P4 or pDONR221-P3P2 for rBiFC and pDONR207 for mbSUS via BP reaction according to the manufacturer’s instructions (Invitrogen). Eventually, LR recombination was performed to clone sequences from the generated entry vectors into the appropriate destination vectors pBiFCt-2in1-NC for rBiFC and pNX35-DEST-1 or pMetOYC-Dest for mbSUS. The resulting final constructs are listed in Table S12. For recombinant protein expression, the coding sequence of NET4A was amplified from Col-0 cDNA and cloned into pET28a(+) (Sigma-Aldrich) using the restriction sites for *Btg*ZI and *Xho*I. Used primers are given in Table S13. All generated constructs were confirmed by sequencing.

### Crossing of Arabidopsis and generation of new lines

For generation of stable Arabidopsis plants, *A. tumefaciens* was transformed as described previously (Jeblick et al., 2023), and floral inoculation performed as described by Nurasaka et al. (Narusaka et al., 2010). The lines lifeact-RFP x NET4A-GFP^OE^, *act7-4* x NET4A-GFP^OE^, NET4A-GFP^OE^ x *amiR-vps16*, SRß-mTurquoise x NET4A-GFP^OE^, RFP-HDEL x NET4A-GFP and RFP-HDEL x NET4A-GFP^OE^ were generated by crossing the respective parental lines as shown in Table S14. Homozygous lines were identified by genotyping using the corresponding genotyping primers in Supplementary Table S15. In case of fluorophore-fusion lines, the offspring of a potential homozygous plant was checked for fluorescence signals to verify homozygosity. CRISPR/Cas9 deletion mutants were additionally selected according to the loss of the CRISPR/Cas9 construct in the following generation. All newly established Arabidopsis lines are homozygous, except two of the crossing lines as stated in Table S14.

### Confocal Microscopy

Live cell imaging, except for rBiFC assays, was performed with a Zeiss LSM880, AxioObserver SP7 confocal laser-scanning microscope, equipped with either a Zeiss C-Apochromat 40×/1.2 W AutoCorr M27 water-immersion objective or a Plan-Apochromat 20x/0.8 M27 objective (INST 248/254-1). Vacuole stainings were carried out as described previously (Scheuring et al., 2015). Cell walls were stained by mounting roots in 0.01 mg/ml PI solution. Fluorescence signals were acquired for mTurquoise (excitation/emission 405 nm/464-482 nm), MDY-64 (excitation/emission 458 nm/473–527 nm), GFP and BCECF (excitation/emission 488 nm/500–571 nm), RFP/mCherry (excitation/emission 594 nm/598–687 nm), FM4-64 (excitation/emission 543 nm/583–758 nm) and PI (excitation/emission 543 nm/580–718 nm), and processed using Zeiss software ZEN 2.3 or Fiji software (https://imagej.net/Fiji). Z-stacks were recorded with a step size of either 400 nm (for quantitative analysis of actin filament organization) or 540 nm (for 3D reconstruction of cells and vacuoles and assessment of vacuolar occupancy). For rBiFC experiments, samples were imaged using the Leica TCS SP8 WLL SMD-FLIM-FCS with an HC PL APO CS2 63x/1.20 (water) objective. Fluorophores were excited using a white laser (514 nm for YFP, 570 nm for RFP) and the emission was detected using hybrid detectors (HyD SMD; 520-560 nm for YFP, 580-630 nm for RFP). For perinuclear ER and nucleus confocal microscopy, *Arabidopsis thaliana* cotyledonae cells were imaged using Zeiss PlanApo x100/1.46 NA oil immersion objective on a Zeiss 880 LSM confocal with Airyscan detector. GFP was excited using a 488 nm laser and emission was detected at 523 nm. RFP was excited using a 561 nm laser and emission was detected at 579 nm. All images were collected in line switch scanning mode with 8x averaging.

### Phenotype Analysis

Root length, calculation of vacuolar morphology index (vac. morph. index, VMI), plasma membrane-tonoplast distance and vacuolar occupancy of the cell (occupancy) were carried out as described previously (Kaiser et al., 2019). VMI measurements of *amiR-vps16* and NET4A-GFP^OE^ x *amiR-vps16* were conducted in the early elongation zone, taken into account cells that were twice as long as wide.

### Quantification of actin filament organization

To investigate the actin filament organization, z-stacks of root epidermal cells were acquired for the late meristematic/early elongated zone. For quantitative analysis, z-stack recordings were processed to half z-stacks, covering early elongated cells of atrichoblast files from the outer cortical confocal section to the cell centre (at the position of the nucleus), and used to create maximum intensity projections. Parallelness of actin filaments was then quantified for a defined rectangular area (24.39×11.62 µm) within a single early elongated atrichoblast cell (excluding cell borders) on each maximum intensity projection using the LPX plugin for ImageJ (<u>URL:</u> https://lpixel.net/services/research/lpixel-imagej-plugins/) according to Higaki (Higaki, 2017). Briefly, a box outlining the defined area of a cell used for quantification was created using the “rectangle” tool of the toolbar menu and added as ROI to the ROI manager. Actin filaments were skeletonized using the “Lpx Filter2d” plugin with “lineFilters” set as filter and “lineExtract” set as linemode. The parameters for “lineExtract” were set to giwsiter = 5, mdnmsLen = 15, pickup = otsu, shaveLen = 5 and delLen = 5. To exclude skeletonized actin filaments outside the desired area of the image, they were masked by inverting the previously created ROI (“Edit→Selection→Make Inverse”) and painting it black using “Edit→Fill” while the intensity value of the “Color picker” tool from the toolbar menu is set to zero. The “Lpx Filter2d” plugin was then used on this masked image with “lineFilters” set as filter and “lineFeature” set as linemode to eventually calculate parallelness (given as “a_normAvgRad” value).

### 3D Surface Rendering

3D reconstruction of cells and vacuoles was performed using Imaris 8.4 (Bitplane) software (https://imaris.oxinst.com/). For the creation of cell models, the manual drawing function (distance) of the surface creation tool was used. Cell borders were marked according to the signals for PI channel on at least every third slice of a z-stack. Subsequently, a surface representing the whole cell was created based on these markings. The surface was then used to produce a masked version of the BCECF-AM channel by using the mask selection function in the edit menu and setting the voxels outside the surface to 0. For the creation of the vacuole models, this masked channel was used to automatically create a second surface corresponding to the BCECF signals. Before completing the surface rendering, the displayed model was visually compared to the underlying BCECF signals and adjustments were made using the absolute intensity threshold option. Eventually, the volumes of the generated 3D models, each given in the statistic menu of the respective surface, were used to calculate the vacuolar occupancy of the cell.

### Protein extraction

For *N. benthamiana* leaf samples from rBIFC experiments, protein extraction was performed using Lyse and Load (LL-) buffer (50 mM Tris/HCl; pH 6.8, including 2 % sodium dodecylsulfate (SDS), 7 M urea, 30 % glycerol, 0.1 M DTT and 0.04 % bromophenol blue. Proteins of cell culture samples from mbSUS assays were extracted using the same LL-buffer according to Asseck and Grefen (Asseck and Grefen, 2018).

### Immunological detection

For immunological detection, samples were separated via sodium dodecylsulfate polyacrylamide gel electrophoresis (SDS–PAGE). If necessary, samples were mixed with loading dye, incubated at 95 °C for 5 min and shortly centrifuged prior to loading. Blotting was performed using either a nitrocellulose (GE Healthcare, USA) or a polyvinylidene difluoride (PVDF) membrane (Sigma-Aldrich). Membranes were blocked with 5 % skim milk powder in TBS-T (150 mM NaCl, 10 mM Tris/HCl pH 8.0, 0.1% Tween 20), washed with TBS-T and then incubated with the first antibody solution. After washing the membrane again, it was probed with the second antibody for 1 h. Subsequently, the membrane was washed another three times at least, submerged with ECL solution (ECL Prime Kit; Amersham, UK) and chemiluminescence signals were detected using the ChemiDoc system from BioRad. The following antibodies were used: monoclonal anti-GFP antibody (1:500; Roche), monoclonal anti-HA antibody (1:1000, Roche), polyclonal anti-VP16 antibody (Genetex; 1:1000), polyclonal anti-NET4 (1:1000).

### Co-Immunoprecipitation and mass spectrometry of peptides

For the co-immunoprecipitation, 7 days old seedlings (Col-0, NET4A-GFP, NET4A-GFP^OE^) were grown on plates. From every plate, 500 mg plant material was harvested, frozen and grinded in liquid nitrogen. Four individual replicates were carried out. 1.5 ml extraction buffer (50 mM NaCl, 50 mM HEPES pH 8, 5 mM MgCl_2_, 1 mM PMSF, 2 mM DSP (crosslinker) and 1x cOmplete^TM^, EDTA-free Protease Inhibitor) was added and the samples incubated for 30 min at 4°C. After quenching, lysates were clarified by centrifugation twice at 16,000 g at 4°C. Supernatant was applied onto GFP-nanobodies coupled to magnetic agarose beads (GFP-trap, ChromoTek, Germany) equilibrated in lysis buffer. Beads were incubated with the lysate on an end over end shaker for 1 h at 4°C. To wash the beads, tubes were magnetized for 1 min, the supernatant discarded and 750 µl wash-buffer (50 mM NaCl, 50 mM Tris-HCL pH 8.0, 0.05% Tween-20; 5 mM MgCl_2_) was added and tubes were agitated on an end over end shaker for 1 min at 4°C. Two wash cycles were done, followed by three in wash-buffer lacking Tween-20. The final bead pellet was resuspended in 100 µl 2xSDS sample buffer (120 mM Tris-HCL pH 6.8, 20% glycerol, 4% SDS, 0.04 bromophenol blue, 100 mM DTT) and boiled for 1 min at 98°C. After magnetizing again, the eluate was transferred into a new tube and 20 µl were loaded on a 10% SDS-PAGE. Samples were run for approximately 1.5 cm into the separating gel, gels were stained with colloidal Coomassie for 3 h. Visible protein bands were cut out and destained for 30 s using destaining solution (10 % acetic acid, 25 % methanol). Before and after cutting, pictures were taken for documentation purposes. After destaining, the cut bands were diced into 1 x 1 mm cubes and transferred into a reaction tube filled with 0.5 ml of 25 % methanol. The prepared samples were then further processed and analyzed by mass spectrometry as described previously (Müller et al., 2018). After analysis, the raw data were processed with MaxQuant (1.6.3.3) using the latest Uniprot Arabidopsis proteome fasta file. Statistical data analysis was performed using Perseus (Tyanova et al., 2016). First, protein LFQ-values were log_2_-transformed, and accessions were removed that were identified in less than two replicates in one of the two categories (Col-0, NET4A-GFP^OE^). For statistics, missing values were imputed, based on all measured log_2_ LFQ intensities, with random numbers. Enrichment between NET4A-GFP^OE^ and Col-0 was determined by one-sided *t*-test and permutation-based FDR of 0.05 and S_0_ = 1 (when both compared sample sets contained at least three valid values).

### Extraction and mass spectrometry analysis of lipids

Root lipids were extracted using approximately 3 mg of dry tissue following the protocol from Gutbrod et al., (Gutbrod et al., 2021). Briefly, phospholipase activity was inactivated by incubating the tissue in boiling water for 5 min. Extraction started by adding 1 ml of chloroform: methanol (1: 2, v/v), followed by 1 ml of chloroform: methanol (2: 1, v/v), and 1 ml of chloroform. For each extraction step, samples were vortexed thoroughly, incubated 5 min and centrifuged and the supernatants collected in a new tube. To the combined supernatant, 0.75 ml of 300 mM ammonium acetate was added, samples were vortexed thoroughly, centrifuged for 5 min at 1500 *g*, and the lower phase transferred to a new tube. Extracts were evaporated under a N_2_ stream and dissolved in mobile phase B [(isopropanol/acetonitrile (90:10, v/v) containing 10 mM ammonium formate and 0.1% (v/v) formic acid].

Lipid extracts were separated on a bio-inert LC-40D Xsi UHPLC system (Shimadzu, Kyoto, Japan) equipped with a Acquity UPLC CSH C18 column (2.1 × 100mm 130Å, 1.7 µm; Waters, Milford, Ma, USA) operated at a 0.6 mL/min flow and equilibrated to 65°C. Mobile phase A consisted of acetonitrile/water (60:40, v/v) containing 10 mM ammonium formate and 0.1% (v/v) formic acid. The gradient was as follows: 15% B at 0 min, increased to 30% B over 2 min, then to 48% B over 0.5 min, followed by an increase to 80% B over 8.5 min. The gradient was further increased to 100% B at 11.5 min and held for 0.5 min, before returning to 15% B for a total time of 15 min. Mass spectrometry analysis was carried out on a ZenoTOF 8600 system and an Optiflow analytical electrospray ionization (ESI) source (both Sciex, Toronto, Canada). Untargeted analysis was performed using the positive ionization mode using information-dependent acquisition (IDA) with a maximum of 25 candidate ions. Dynamic background subtraction was enabled and precursor ions were excluded after being selected twice. Collision induced dissociation was employed with a collision energy spread of 5 V. See Suppl. Dataset 1 for detailed information. Data was processed and annotated using MS-DIAL software (v5.5) (Tsugawa et al., 2015). Mass accuracy tolerances were set to 0.01 Da for MS1 and 0.025 Da for MS2. The peak height was set to a minimum of 500 amplitude. Lipid annotation used an LBM database (Msp20250929174203_NCDK_conventional_converted_dev) (Tsugawa et al., 2020). For the alignment, the retention time tolerance was set to 0.17 min, and MS1 tolerance to 0.015 Da. After identification by the software, the spectra of the lipid species depicted in Figure 8 were manually confirmed.

### Ratiometric Bimolecular Fluorescence Complementation (rBiFC)

Agrobacterium-mediated transient transformation of approximately 4-week old *N. benthamiana* leaves with 2in1 rBiFC constructs was performed as described previously (Mehlhorn et al., 2018). Fluorescence intensities were recorded 2 days’ post infiltration for approximately 20-30 images per construct. Ratios between complemented YFP fluorescence and RFP were calculated and plotted using Fiji and Graphpad Prism5 software. Immunoblotting verified protein expression utilizing anti-NET4A antibody and anti-HA peroxidase conjugated antibody, respectively.

### Mating-based Split-Ubiquitin System (mbSUS)

Bait/Cub fusions and prey/Nub fusions were generated by cloning the desired genes of interest into the Gateway-compatible vectors pMetOYC-Dest and pNX35-Dest, respectively. These were transformed into haploid yeast strains THY.AP4 (Cub fusion) and THY.AP5 (Nub fusions) as described previously (Asseck and Grefen, 2018). After mating, diploid yeasts were dropped in 10 times OD dilutions on growth control (CSM-Leu-, Trp-, Ura-) and interaction-selective media (CSM-Leu-, Trp-, Ura-, Ade-, His-) supplemented with 50 or 500 μM methionine. The non-mutated N-terminal ubiquitin moiety (NubI) was used as positive control, and NubI13G (NubG) as negative control. Protein expression was verified in haploid yeast using antibodies against the VP16 domain for the Cub fusion or the HA epitope tag for the Nub fusions.

### Transient gene expression

Transient expression was performed as previously described (Jeblick et al., 2023). Infiltrated areas were marked for easier identification and plants were incubated under reduced light conditions for 48-72 h prior microscopic analysis. For constructs with estradiol-inducible promoter, 24 h post infiltration the abaxial leaf sides of infiltrated areas were painted with 100 µM β-estradiol solution. Plants were then further incubated for additional 3-4 days prior analysis.

### Gene Accession Codes

Sequence data of gene and protein sequences used in this article can be found at the Arabidopsis Information Resource (TAIR; http://www.arabidopsis.org/), GenBank/EMBL, Aramemnon (http://aramemnon.uni-koeln.de/), or UniProt (https://www.uniprot.org/proteomes/UP000006548databases) under the following accession numbers: NET4A (At5g58320), NET4B (At2g30500), NET3A (At1g03470), NET3C (At2g47920), RABG3A (At4g09720), RABG3B (At1g22740, VPS16 (At2g38020), VAMP711 (At4g32150), RABC1 (At1g43890), VAP27-1 (At3g60600), ORP1C (At4g08180), ORP2A (At4g22540) and ORP3C (At5g59420).

### Statistical analysis

All quantitative data was analyzed using the GraphPad Prism 5 or 9 software (https://www.graphpad.com/scientific-software/prism/). The precise statistical method used is given in the respective Figure legends. Box-plot borders in the graphs represent 25th to 75th percentile, the horizontal line the median and whiskers minimum to maximum values.

## Data, Materials, and Software Availability

Study data are presented in the article or supporting information.

## Acknowledgements

We would like to thank Takashi Ueda for providing the *rabg3a,b,c,d,e,f* sextuple mutant and Joe McKenna for the RFP-HDEL Arabidopsis line and plasmid. We thank Matthias Hahn for critical reading of the manuscript and constructive advice throughout the project. Technical help from Patrick Pattar, Anne Lau, Sophie Eisele, Benjamin Ziehmer and Tom Niehoff is highly appreciated. For the kind willingness to share facilities and equipment, we thank Ekkehard Neuhaus. This work was supported by grants from the *BioComp* research initiative (Rhineland-Palatinate, Germany) and the German research foundation (DFG; SCHE 1836/4-1 and SCHE 1836/4-2) to D.S.

## Author contributions

D.S. conceived and outlined the research. S.K. planned and performed most experiments and created the majority of constructs and Arabidopsis lines. D.M. and C.G. helped with planning and creation of rBiFC and mbSUS constructs and performed rBiFC and part of mbSUS experiments. P.R.M. performed part of the analysis of *net3 net4* triple mutants. F.R. and F.W. helped with Co-IPs and statistical evaluation of MS/MS data. F.S. and M.S. helped with planning and performed LC-MS/MS analysis. N.F. and V.K. performed microscopy for colocalization studies of RFP-HDEL and NET4A-GFP in Arabidopsis. A.CV. and T.I. carried out and analysed for lipidomics. K.S. provided the *amiR-vps16* line. S.K. and D.S. evaluated data, assembled Figures and wrote the manuscript. All co-authors saw and commented on the manuscript.

## Competing interests

The authors declare no competing interests.

## Notes

### Competing Interest Statement

The authors have declared no competing interest.

### Summary of Updates

This manuscript has been extensively revised, including its title, abstract, organization of the Results, and Discussion. The revised version incorporates additional experimental data, updated figures and supplementary information, and a refined mechanistic model. Particular emphasis is placed on the overlapping functions of NET3 and NET4 proteins in actin-dependent vacuolar organization and their association with the endoplasmic reticulum.

## References

Aniento, F., Sánchez de Medina Hernández, V., Dagdas, Y., Rojas-Pierce, M., and Russinova, E. (2022). Molecular mechanisms of endomembrane trafficking in plants. The Plant Cell 34 (1): 146–173.

Asseck, L.Y., and Grefen, C. (2018). Detecting Interactions of Membrane Proteins: The Split-Ubiquitin System. Methods in molecular biology (Clifton, N.J.) 1794: 49–60.

Balderhaar, H.J.k., and Ungermann, C. (2013). CORVET and HOPS tethering complexes - coordinators of endosome and lysosome fusion. Journal of cell science 126 (Pt 6): 1307–1316.

Brillada, C., Zheng, J., Krüger, F., Rovira-Diaz, E., Askani, J.C., Schumacher, K., and Rojas-Pierce, M. (2018). Phosphoinositides control the localization of HOPS subunit VPS41, which together with VPS33 mediates vacuole fusion in plants. Proceedings of the National Academy of Sciences of the United States of America 115 (35): E8305–E8314.

Cabantous, S., Terwilliger, T.C., and Waldo, G.S. (2005). Protein tagging and detection with engineered self-assembling fragments of green fluorescent protein. Nature biotechnology 23 (1): 102–107.

Chytilova, E., Macas, J., Sliwinska, E., Rafelski, S.M., Lambert, G.M., and Galbraith, D.W. (2000). Nuclear dynamics in Arabidopsis thaliana. Molecular Biology of the Cell 11 (8): 2733–2741.

Cui, Y., Zhao, Q., Hu, S., and Jiang, L. (2020). Vacuole Biogenesis in Plants: How Many Vacuoles, How Many Models? Trends in plant science 25 (6): 538–548.

Deeks, M.J., Calcutt, J.R., Ingle, E.K.S., Hawkins, T.J., Chapman, S., Richardson, A.C., Mentlak, D.A., Dixon, M.R., Cartwright, F., Smertenko, A.P., Oparka, K., and Hussey, P.J. (2012). A superfamily of actin-binding proteins at the actin-membrane nexus of higher plants. Current biology CB 22 (17): 1595–1600.

Denninger, P., Reichelt, A., Schmidt, V.A.F., Mehlhorn, D.G., Asseck, L.Y., Stanley, C.E., Keinath, N.F., Evers, J.-F., Grefen, C., and Grossmann, G. (2019). Distinct RopGEFs Successively Drive Polarization and Outgrowth of Root Hairs. Current biology CB 29 (11): 1854–1865.e5.

Duckney, P.J., Wang, P., and Hussey, P.J. (2022). Membrane contact sites and cytoskeleton-membrane interactions in autophagy. FEBS letters 596 (17): 2093–2103.

Ebine, K., Inoue, T., Ito, J., Ito, E., Uemura, T., Goh, T., Abe, H., Sato, K., Nakano, A., and Ueda, T. (2014). Plant vacuolar trafficking occurs through distinctly regulated pathways. Current biology CB 24 (12): 1375–1382.

Eitzen, G., Wang, L., Thorngren, N., and Wickner, W. (2002). Remodeling of organelle-bound actin is required for yeast vacuole fusion. The Journal of cell biology 158 (4): 669–679.

Geldner, N., Dénervaud-Tendon, V., Hyman, D.L., Mayer, U., Stierhof, Y.-D., and Chory, J. (2009). Rapid, combinatorial analysis of membrane compartments in intact plants with a multicolor marker set. The Plant journal for cell and molecular biology 59 (1): 169–178.

Gilliland, L.U., Pawloski, L.C., Kandasamy, M.K., and Meagher, R.B. (2003). Arabidopsis actin gene ACT7 plays an essential role in germination and root growth. The Plant journal for cell and molecular biology 33 (2): 319–328.

Grefen, C., Obrdlik, P., and Harter, K. (2009). The determination of protein-protein interactions by the mating-based split-ubiquitin system (mbSUS). Methods in molecular biology (Clifton, N.J.) 479: 217–233.

Gundersen, G.G., and Worman, H.J. (2013). Nuclear positioning. Cell 152 (6): 1376–1389.

Gutbrod, K., Peisker, H., and Dörmann, P. (2021). Direct Infusion Mass Spectrometry for Complex Lipid Analysis. Methods in molecular biology (Clifton, N.J.) 2295: 101–115.

Hanaoka, K., Nishikawa, K., Ikeda, A., Schlarmann, P., Sasaki, S., Fujii, S., Yamashita, S., Nakaji, A., and Funato, K. (2024). Membrane contact sites regulate vacuolar fission via sphingolipid metabolism. eLife 12.

Hawkins, T.J., Kopischke, M., Duckney, P.J., Rybak, K., Mentlak, D.A., Kroon, J.T.M., Bui, M.T., Richardson, A.C., Casey, M., Alexander, A., Jaeger, G. de, Kalde, M., Moore, I., Dagdas, Y., Hussey, P.J., and Robatzek, S. (2023). NET4 and RabG3 link actin to the tonoplast and facilitate cytoskeletal remodelling during stomatal immunity. Nature communications 14 (1): 5848.

Higaki, T. (2017). Quantitative evaluation of cytoskeletal organizations by microscopic image analysis. PLANT MORPHOL 29 (1): 15–21.

Jeblick, T., Leisen, T., Steidele, C.E., Albert, I., Müller, J., Kaiser, S., Mahler, F., Sommer, F., Keller, S., Hückelhoven, R., Hahn, M., and Scheuring, D. (2023). Botrytis hypersensitive response inducing protein 1 triggers noncanonical PTI to induce plant cell death. Plant Physiology 191 (1): 125–141.

Kader, J.-C. (1996). LIPID-TRANSFER PROTEINS IN PLANTS. Annual review of plant physiology and plant molecular biology 47: 627–654.

Kaiser, S., Eisa, A., Kleine-Vehn, J., and Scheuring, D. (2019). NET4 Modulates the Compactness of Vacuoles in Arabidopsis thaliana. International journal of molecular sciences 20 (19).

Kaiser, S., Eisele, S., and Scheuring, D. (2021). Vacuolar occupancy is crucial for cell elongation and growth regardless of the underlying mechanism. Plant Signaling & Behavior 16 (8): 1922796.

Kaiser, S., and Scheuring, D. (2020). To Lead or to Follow: Contribution of the Plant Vacuole to Cell Growth. Frontiers in Plant Science 11: 553.

Kandasamy, M.K., McKinney, E.C., and Meagher, R.B. (2009). A single vegetative actin isovariant overexpressed under the control of multiple regulatory sequences is sufficient for normal Arabidopsis development. The Plant Cell 21 (3): 701–718.

Ketelaar, T., Faivre-Moskalenko, C., Esseling, J.J., Ruijter, N.C.A. de, Grierson, C.S., Dogterom, M., and Emons, A.M.C. (2002). Positioning of nuclei in Arabidopsis root hairs: an actin-regulated process of tip growth. The Plant Cell 14 (11): 2941–2955.

Kost, B., and Chua, N.-H. (2002). The plant cytoskeleton: vacuoles and cell walls make the difference. Cell 108 (1): 9–12.

Labun, K., Montague, T.G., Gagnon, J.A., Thyme, S.B., and Valen, E. (2016). CHOPCHOP v2: a web tool for the next generation of CRISPR genome engineering. Nucleic Acids Research 44 (W1): W272–6.

Lampropoulos, A., Sutikovic, Z., Wenzl, C., Maegele, I., Lohmann, J.U., and Forner, J. (2013). GreenGate - A Novel, Versatile, and Efficient Cloning System for Plant Transgenesis. PLOS ONE 8 (12).

Lupanga, U., Röhrich, R., Askani, J., Hilmer, S., Kiefer, C., Krebs, M., Kanazawa, T., Ueda, T., and Schumacher, K. (2020). The Arabidopsis V-ATPase is localized to the TGN/EE via a seed plant-specific motif. eLife 9.

McDowell, J.M., An, Y.Q., Huang, S., McKinney, E.C., and Meagher, R.B. (1996). The arabidopsis ACT7 actin gene is expressed in rapidly developing tissues and responds to several external stimuli. Plant Physiology 111 (3): 699–711.

Mehlhorn, D.G., Wallmeroth, N., Berendzen, K.W., and Grefen, C. (2018). 2in1 Vectors Improve In Planta BiFC and FRET Analyses. Methods in molecular biology (Clifton, N.J.) 1691: 139–158.

Müller, J., König, Y., Xhelilaj, K., Kaiser, S., Müller, T., Vielba-Fernández, A., Gronnier, J., Löfke, C., Krebs, M., and Scheuring, D. (2025). Salicylic acid inhibits V-ATPase activity and restricts cell elongation. Plant Physiology 199 (2).

Müller, N., Leroch, M., Schumacher, J., Zimmer, D., Könnel, A., Klug, K., Leisen, T., Scheuring, D., Sommer, F., Mühlhaus, T., Schroda, M., and Hahn, M. (2018). Investigations on VELVET regulatory mutants confirm the role of host tissue acidification and secretion of proteins in the pathogenesis of Botrytis cinerea. The New phytologist 219 (3): 1062–1074.

Narusaka, M., Shiraishi, T., Iwabuchi, M., and Narusaka, Y. (2010). The floral inoculating protocol: a simplified Arabidopsis thaliana transformation method modified from floral dipping. Plant Biotechnology 27 (4): 349–351.

Pfaffl, M.W. (2001). A new mathematical model for relative quantification in real-time RT-PCR. Nucleic Acids Research 29 (9): e45.

Renna, L., Stefano, G., Puggioni, M.P., Kim, S.-J., Lavell, A., Froehlich, J.E., Burkart, G., Mancuso, S., Benning, C., and Brandizzi, F. (2024). ER-associated VAP27-1 and VAP27-3 proteins functionally link the lipid-binding ORP2A at the ER-chloroplast contact sites. Nature communications 15 (1): 6008.

Riedl, J., Crevenna, A.H., Kessenbrock, K., Yu, J.H., Neukirchen, D., Bista, M., Bradke, F., Jenne, D., Holak, T.A., Werb, Z., Sixt, M., and Wedlich-Soldner, R. (2008). Lifeact: a versatile marker to visualize F-actin. Nature Methods 5 (7): 605–607.

Rojo, E., Gillmor, C.S., Kovaleva, V., Somerville, C.R., and Raikhel, N.V. (2001). VACUOLELESS1 is an essential gene required for vacuole formation and morphogenesis in Arabidopsis. Developmental cell 1 (2): 303–310.

Rosero, A., Oulehlová, D., Stillerová, L., Schiebertová, P., Grunt, M., Žárský, V., and Cvrčková, F. (2016). Arabidopsis FH1 Formin Affects Cotyledon Pavement Cell Shape by Modulating Cytoskeleton Dynamics. Plant & cell physiology 57 (3): 488–504.

Scheuring, D., Löfke, C., Krüger, F., Kittelmann, M., Eisa, A., Hughes, L., Smith, R.S., Hawes, C., Schumacher, K., and Kleine-Vehn, J. (2016). Actin-dependent vacuolar occupancy of the cell determines auxin-induced growth repression. Proceedings of the National Academy of Sciences of the United States of America 113 (2): 452–457.

Scheuring, D., Minina, E.A., Krueger, F., Lupanga, U., Krebs, M., and Schumacher, K. (2024). Light at the end of the tunnel: FRAP assays combined with super resolution microscopy confirm the presence of a tubular vacuole network in meristematic plant cells. The Plant Cell 36 (11): 4683–4691.

Scheuring, D., Schöller, M., Kleine-Vehn, J., and Löfke, C. (2015). Vacuolar staining methods in plant cells. Methods in molecular biology (Clifton, N.J.) 1242: 83–92.

Scorrano, L., Matteis, M.A. de, Emr, S., Giordano, F., Hajnóczky, G., Kornmann, B., Lackner, L.L., Levine, T.P., Pellegrini, L., Reinisch, K., Rizzuto, R., Simmen, T., Stenmark, H., Ungermann, C., and Schuldiner, M. (2019). Coming together to define membrane contact sites. Nature communications 10 (1): 1287.

Shvarev, D., Schoppe, J., König, C., Perz, A., Füllbrunn, N., Kiontke, S., Langemeyer, L., Januliene, D., Schnelle, K., Kümmel, D., Fröhlich, F., Moeller, A., and Ungermann, C. (2022). Structure of the HOPS tethering complex, a lysosomal membrane fusion machinery. eLife 11.

Starr, M.L., and Fratti, R.A. (2019). The Participation of Regulatory Lipids in Vacuole Homotypic Fusion. Trends in biochemical sciences 44 (6): 546–554.

Stemmer, M., Thumberger, T., Del Sol Keyer, M., Wittbrodt, J., and Mateo, J.L. (2015). CCTop: An Intuitive, Flexible and Reliable CRISPR/Cas9 Target Prediction Tool. PLOS ONE 10 (4): e0124633.

Stephani, M., Picchianti, L., Gajic, A., Beveridge, R., Skarwan, E., Sanchez de Medina Hernandez, V., Mohseni, A., Clavel, M., Zeng, Y., Naumann, C., Matuszkiewicz, M., Turco, E., Loefke, C., Li, B., Dürnberger, G., Schutzbier, M., Chen, H.T., Abdrakhmanov, A., Savova, A., Chia, K.-S., Djamei, A., Schaffner, I., Abel, S., Jiang, L., Mechtler, K., Ikeda, F., Martens, S., Clausen, T., and Dagdas, Y. (2020). A cross-kingdom conserved ER-phagy receptor maintains endoplasmic reticulum homeostasis during stress. eLife 9.

Stührwohldt, N., Scholl, S., Lang, L., Katzenberger, J., Schumacher, K., and Schaller, A. (2020). The biogenesis of CLEL peptides involves several processing events in consecutive compartments of the secretory pathway. eLife 9.

Takemoto, K., Ebine, K., Askani, J.C., Krüger, F., Gonzalez, Z.A., Ito, E., Goh, T., Schumacher, K., Nakano, A., and Ueda, T. (2018). Distinct sets of tethering complexes, SNARE complexes, and Rab GTPases mediate membrane fusion at the vacuole in Arabidopsis. Proceedings of the National Academy of Sciences of the United States of America 115 (10): E2457–E2466.

Tan, J.X., and Finkel, T. (2022). A phosphoinositide signalling pathway mediates rapid lysosomal repair. Nature 609 (7928): 815–821.

Tsugawa, H., Cajka, T., Kind, T., Ma, Y., Higgins, B., Ikeda, K., Kanazawa, M., VanderGheynst, J., Fiehn, O., and Arita, M. (2015). MS-DIAL: data-independent MS/MS deconvolution for comprehensive metabolome analysis. Nat Methods 12 (6): 523–526.

Tsugawa, H., Ikeda, K., Takahashi, M., Satoh, A., Mori, Y., Uchino, H., Okahashi, N., Yamada, Y., Tada, I., Bonini, P., Higashi, Y., Okazaki, Y., Zhou, Z., Zhu, Z.-J., Koelmel, J., Cajka, T., Fiehn, O., Saito, K., Arita, M., and Arita, M. (2020). A lipidome atlas in MS-DIAL 4. Nat Biotechnol 38 (10): 1159–1163.

Tyanova, S., Temu, T., and Cox, J. (2016). The MaxQuant computational platform for mass spectrometry-based shotgun proteomics. Nature protocols 11 (12): 2301–2319.

Vale, R.D. (2003). The molecular motor toolbox for intracellular transport. Cell 112 (4): 467–480.

Viotti, C., Krüger, F., Krebs, M., Neubert, C., Fink, F., Lupanga, U., Scheuring, D., Boutté, Y., Frescatada-Rosa, M., Wolfenstetter, S., Sauer, N., Hillmer, S., Grebe, M., and Schumacher, K. (2013). The endoplasmic reticulum is the main membrane source for biogenesis of the lytic vacuole in Arabidopsis. The Plant Cell 25 (9): 3434–3449.

Waadt, R., Krebs, M., Kudla, J., and Schumacher, K. (2017). Multiparameter imaging of calcium and abscisic acid and high-resolution quantitative calcium measurements using R-GECO1-mTurquoise in Arabidopsis. New Phytol 216 (1): 303–320.

Wang, P., Duckney, P., Gao, E., Hussey, P.J., Kriechbaumer, V., Li, C., Zang, J., and Zhang, T. (2023). Keep in contact: multiple roles of endoplasmic reticulum-membrane contact sites and the organelle interaction network in plants. The New phytologist 238 (2): 482–499.

Wang, P., Hawkins, T.J., Richardson, C., Cummins, I., Deeks, M.J., Sparkes, I., Hawes, C., and Hussey, P.J. (2014). The plant cytoskeleton, NET3C, and VAP27 mediate the link between the plasma membrane and endoplasmic reticulum. Current biology CB 24 (12): 1397–1405.

Wang, Z.-P., Xing, H.-L., Dong, L., Zhang, H.-Y., Han, C.-Y., Wang, X.-C., and Chen, Q.-J. (2015). Egg cell-specific promoter-controlled CRISPR/Cas9 efficiently generates homozygous mutants for multiple target genes in Arabidopsis in a single generation. Genome biology 16 (1): 144.

Wickner, W. (2010). Membrane fusion: five lipids, four SNAREs, three chaperones, two nucleotides, and a Rab, all dancing in a ring on yeast vacuoles. Annual review of cell and developmental biology 26: 115–136.

Xing, H.-L., Dong, L., Wang, Z.-P., Zhang, H.-Y., Han, C.-Y., Liu, B., Wang, X.-C., and Chen, Q.-J. (2014). A CRISPR/Cas9 toolkit for multiplex genome editing in plants. BMC plant biology 14: 327.

Ye, H., Gao, J., Liang, Z., Lin, Y., Yu, Q., Huang, S., and Jiang, L. (2022). Arabidopsis ORP2A mediates ER-autophagosomal membrane contact sites and regulates PI3P in plant autophagy. Proceedings of the National Academy of Sciences of the United States of America 119 (43): e2205314119.

Zang, J., Klemm, S., Pain, C., Duckney, P., Bao, Z., Stamm, G., Kriechbaumer, V., Bürstenbinder, K., Hussey, P.J., and Wang, P. (2021). A novel plant actin-microtubule bridging complex regulates cytoskeletal and ER structure at ER-PM contact sites. Current biology CB 31 (6): 1251–1260.e4.

Zhao, Y., Liu, Z., Wang, L., and Liu, H. (2022). Fumonisin B1 as a Tool to Explore Sphingolipid Roles in Arabidopsis Primary Root Development. International journal of molecular sciences 23 (21).

Zhou, W.-J., Li, H., Zhang, K.-K., Wang, F., Chu, X., and Jiang, J.-H. (2021). Genetically Encoded Sensor Enables Endogenous RNA Imaging with Conformation-Switching Induced Fluorogenic Proteins. Journal of the American Chemical Society 143 (35): 14394–14401.

