## Supplemental tables for "NETWORKED proteins link actin-dependent vacuolar organization to the endoplasmic reticulum"

### Supplementary Tables

**Table S1. Proteins identified by LC-MS/MS after co-immunoprecipitation with NET4A-GFP.** Listed are all candidates that were significantly enriched compared to the WT control. Statistical analysis was based on LFQ intensities and performed using Perseus software. Respective LFQ intensities per sample are shown. Interaction candidates used for further investigation are highlighted in green.

| Protein names | Gene names | LFQ intensities | | | | | | | | | | | |
|---|---|---|---|---|---|---|---|---|---|---|---|---|---|
| | | WT | | | | NET4A-GFP | | | | NET4A-GFP$^{OE}$ | | | |
| | | #1 | #2 | #3 | #4 | #5 | #6 | #7 | #8 | #9 | #10 | #11 | #12 |
| Protein NETWORKED 4A | *NET4A* | 0 | 0 | 0 | 0 | 0 | 6381.9 | 7097.7 | 9505.9 | 1374900 | 1765600 | 538860 | 785530 |
| Ras-related protein RABG3a | *RABG3A* | 0 | 0 | 0 | 0 | 1741.8 | 589.66 | 1820.1 | 694.31 | 60257 | 33834 | 12842 | 15608 |
| | At4g01245 | 0 | 0 | 0 | 0 | 0 | 0 | 0 | 0 | 9929.8 | 12561 | 4714.3 | 9260 |
| Ras-related protein RABC1 | *RABC1* | 0 | 0 | 0 | 0 | 0 | 0 | 0 | 0 | 5426.8 | 3518 | 2908.1 | 1899.3 |
| Ubiquitin-40S ribosomal protein S27a-1/-2/-3; Ubiquitin-60S ribosomal protein L40-1/-2; Ubiquitin-NEDD8-like protein RUB1/2; Polyubiquitin 3/4/8/9/10/11/12/14; Ubiquitin-related 1/2/3/4/5/6/7/8; | *RPS27AA/AB/AC; RPL40A/B; RUB1/2; UBQ3/4/8/9/10/11/12/13/14;* | 0 | 0 | 2926.3 | 1222.1 | 0 | 2078.7 | 2902.1 | 2784.7 | 38306 | 23426 | 15116 | 6464.5 |
| | At2g15042 | 0 | 0 | 0 | 0 | 0 | 0 | 0 | 0 | 5692.6 | 0 | 4714.3 | 9260 |
| Probable mediator of RNA polymerase II transcription subunit 37e/-c; Heat shock 70 kDa protein 18 | *MED37E; HSC70-1; MED37D; HSP70-18* | 2011.7 | 3523.9 | 9823.7 | 8344.5 | 0 | 3462.9 | 6974.1 | 9091.2 | 9279.2 | 49267 | 23070 | 34804 |
| Ras-related protein RABG3b | *RABG3B* | 0 | 0 | 0 | 0 | 0 | 1084.3 | 0 | 0 | 14639 | 13946 | 5311.3 | 0 |
| Heat shock 70 kDa protein 3 | *HSP70-3* | 2323 | 3811.3 | 10049 | 8685.8 | 0 | 3539.9 | 7407.2 | 8653.8 | 5843.1 | 41020 | 15908 | 28887 |
| Probable mediator of RNA polymerase II transcription subunit 37c | *MED37C* | 2358.3 | 3969.2 | 10441 | 8836.1 | 0 | 3605.1 | 7493.6 | 8993 | 5938.1 | 41256 | 16000 | 29054 |

**Table S2. Primers used for qRT-PCR.**

| Primer name | Primer sequence | Gene to detect |
|---|---|---|
| qRT_NET4A_fw | GCTGGAAGCCAATGTGCGTTATC | *NET4A* |
| qRT_NET4A_rev | CGACTTATCTCCGACTCTAGCTCACTC | |
| qRT_NET4B_fw | CGTCTACGGCTCAGAGCAAG | *NET4B* |
| qRT_NET4B_rev | GCTGGATTAACTCGGGACGTTTC | |
| qRT_PP2A_fw | TCGTGCAGTATCGCTTCTCG | *PP2A* (housekeeping) |
| qRT_PP2A_rev | CACAACCGCTTGGTCGACT | |

**Table S3. List of used expression constructs generated by GreenGate cloning.** For all constructs, the destination vector pGGZ003 was used as final Module Z. Entry vectors for Module A to F are given for each construct. For vectors harboring two expression cassettes, single cassettes were assembled separately using intermediate vector pGGM000 together with FH adapter pGGG004 for Module F and intermediate vector pGGN000 together with HA adapter pGGG005 as an additional Module, before combining both cassettes into pGGZ003.

| **Expression construct** | **Module A (promoter)** | **Module B (N-terminal tag)** | **Module C (coding sequence)** | **Module D (C-terminal tag)** | **Module E (terminator)** | **Module F (selection marker)** |
|---|---|---|---|---|---|---|
| NET4B-mCherry$^{OE}$ | pGGA004 | pGGB003 | pGGC-NET4B | pGGD010 | pGGE001 | pGGF Alli-mCherry |
| NET4B:: NET4B-GFP | NET4B promoter (PCR-product) | pGGB003 | pGGC-NET4B | pGGD011 | pGGE001 | pGGF Alli-mCherry |
| NET4A$^{1\text{-}105}$-GFP | pGGA004 | pGGB003 | pGGC-NET4A$^{1\text{-}105}$ | pGGD011 | pGGE001 | pGGF Alli-mCherry |
| NET4A$^{106\text{-}503}$-GFP | pGGA004 | pGGB003 | pGGC-NET4A$^{106\text{-}503}$ | pGGD011 | pGGE001 | pGGF Alli-mCherry |
| NET4A$^{504\text{-}542}$-GFP | pGGA004 | pGGB003 | pGGC-NET4A$^{504\text{-}542}$ | pGGD011 | pGGE001 | pGGF Alli-mCherry |
| NET4A$^{504\text{-}558}$-GFP | pGGA004 | pGGB003 | pGGC-NET4A$^{504\text{-}558}$ | pGGD011 | pGGE001 | pGGF Alli-mCherry |
| NET4A-mTurquoise | pGGA006 | pGGB003 | pGGC-NET4A | pGGD-GSL-mTurquoise | pGGE001 | pGGF Alli-mCherry |
| NET4A::GUS-GFP | pGGA-NET4A | pGGB003 | pGGC051 | pGGD011 | pGGE001 | pGGF Alli-mCherry |
| NET4B::GUS-GFP | NET4B promoter (PCR-product) | pGGB003 | pGGC051 | pGGD011 | pGGE001 | pGGF Alli-mCherry |
| NET4B:: NET4B-mCherry | NET4B promoter (PCR-product) | pGGB003 | pGGC-NET4B | pGGD010 | pGGE001 | pGGF Alli-YFP |
| GFP1-10-VAP27-1// NET4A-GFP11 | pGGA006// pGGA-Ubi-XVE_oLexA-35S | pGGB-GFP1-10// pGGB003 | pGGC-VAP27-1// pGGC-NET4A | pGGD002// pGGD-GFP11 | pGGE001// pGGE001 | -// pGGF Alli-mCherry |
| GFP1-10-VAP27-1// NET4A-GFP11x2 | pGGA006// pGGA-Ubi-XVE_oLexA-35S | pGGB-GFP1-10// pGGB003 | pGGC-VAP27-1// pGGC-NET4A | pGGD002// pGGD-GFP11x2 | pGGE001// pGGE001 | -// pGGF Alli-mCherry |

**Table S4. Primers used for GreenGate cloning.** Bases introducing mutations are displayed in red.

| Primer name | Primer sequence | Purpose |
|---|---|---|
| ggNET4B_fw | AACAGGTCTCAGGCTCAACAATGGCTTCGTCTACGGCTCAG | Amplification of *NET4B* coding sequence |
| ggNET4B_rev | AACAGGTCTCTCTGAAGTTGATAAGACCACTACTCTCTTACTCTTATGTCC | |
| ggNET4A1/2_FMP | GGGATAGTCATATCGGACTCAAAAATTCTAAGTGGC | mutating internal *Bsa*I site in *NET4A* coding sequence |
| ggNET4A1/2_RMP | GCCACTTAGAATTTTTGAGTCCGATATGACTATCCC | |
| ggNET4A1/2_fw | AACAGGTCTCAGGCTCAACAATGGATTATGATCTGCTTCGTTCCAAGAAG | Amplification of full and partial *NET4A* coding sequences |
| ggNET4A-NAB_rev | AACAGGTCTCTCTGAAGAGCCTTTTCTCAACTCTCCGG | |
| ggNET4A1-tr_fw | AACAGGTCTCAGGCTCAACAATGCCTTTAGAGCTTCAGTCACAGGG | |
| ggNET4A-tr_rev | AACAGGTCTCTCTGACTTCTCCATCTCCATAGTCCTATCGTC | |
| ggNET4A-IRQ_fw | AACAGGTCTCAGGCTCAACAATGGAGGTGGAGAAGCAGAGGAGG | |
| ggNET4A-IRQ_rev | AACAGGTCTCTCTGATATGCGTAGTCTCTTGTATTCGTCTCTGG | |
| ggNET4A1_rev | AACAGGTCTCTCTGAAGAAGCAAGAATGGATGATGGTCTTGTTG | |
| gg-pNET4A1_fw | AACAGGTCTCAACCTAGGCTCGTCGTAGTCATCCAAAG | Amplification of 1900bp upstream of *NET4A* start codon (promoter region) |
| gg-pNET4A1_rev | AACAGGTCTCTTGTTGGCTGCAAAAATCAATGGACCCTATG | |
| Esp3I-pNET4B_fw | AACACGTCTCAACCTTGCGGTCTCGAAAAAGTTATAGAGG | Amplification of 1900bp upstream of *NET4B* start codon (promoter region) |
| Esp3I-pNET4B_rev | AACACGTCTCTTGTTGACCGTGGAGTGCAGAG | |
| ggVAP27-1_fw | AACAGGTCTCAGGCTCAACAATGAGTAACATCGATCTGATTGG | Amplification of *VAP27-1* coding sequence |
| ggVAP27-1_rev | AACAGGTCTCTCTGATGTCCTCTTCATAATGTATCCCA | |
| ggGFP1-10-Ntag_fw | AACAGGTCTCAAACAATGTCCAAAGGAGAAGAACTGTTTACCG | Amplification of *GFP1-10, GFP11 and GFP11x2* from plasmid GFP1-10-miniCMV-GFP11×11-GFP-tDeg (Addgene plasmid # 185404 from Jian-Hui Jiang; Zhou et al., 2021) |
| ggGFP1-10-Ntag_rev | AACAGGTCTCTAGCCTTTTTCATTTGGATCTTTGCTCAGGACTG | |
| ggGFP11-Ctag_fw | AACAGGTCTCATCAGGTATGCGTGACCACATGGTCCTTC | |
| ggGFP11-Ctag_rev | AACAGGTCTCTGCAGCTATGTAATCCCAGCAGCATTTACATACTC | |
| ggGFP11-Ctag_rev2 | AACAGGTCTCTGCAGCTAAGTGATGCCTGCGGCG | |

**Table S5. List of entry vectors used for GreenGate cloning.**

| Plasmid | Description | Source |
|---|---|---|
| pGGA000 | Empty entry vector for promoter | Lampropoulos, et al., 2013 |
| pGGA004 | 35S (Cauliflower mosaic virus) promoter | Lampropoulos, et al., 2013 |
| pGGA006 | UBQ10 (UBIQUITIN10) promoter | Lampropoulos, et al., 2013 |
| pGGA-Ubi-XVE_oLexA-35S | Estradiol-inducible promoter | Denninger, et al., 2019 |
| pGGA-NET4A | NET4A (NETWORKED 4A) promoter; 1900bp upsteam of start codon | This work |
| pGGB000 | Empty entry vector for N-terminal tag | Lampropoulos, et al., 2013 |
| pGGB003 | B-dummy (short default random sequence) | Lampropoulos, et al., 2013 |
| pGGB-GFP1-10 | ß-strand 1 to 10 of split GFP (GFP1-10) | This work |
| pGGC000 | Empty entry vector for coding sequence | Lampropoulos, et al., 2013 |
| pGGC051 | GUS (*E. coli* ß-GLUCURONIDASE) | Lampropoulos, et al., 2013 |
| pGGC-NET4B | NET4B (NETWORKED 4B) coding sequence | This work |
| pGGC-NET4A | NET4A (NETWORKED 4A) coding sequence | This work |
| pGGC-NET4A$^{1-105}$ | Partial NET4A coding sequence, aa 1-105 | This work |
| pGGC-NET4A$^{106-503}$ | Partial NET4A coding sequence, aa 106-503 | This work |
| pGGC-NET4A$^{504-542}$ | Partial NET4A coding sequence, aa 504-542 | This work |
| pGGC-NET4A$^{504-558}$ | Partial NET4A coding sequence, aa 504-558 | This work |
| pGGC-VAP27-1 | VAP27-1 coding sequence | This work |
| pGGD000 | Empty entry vector for C-terminal tag | Lampropoulos, et al., 2013 |
| pGGD002 | D-dummy (short default random sequence) | Lampropoulos, et al., 2013 |
| pGGD010 | mCherry | Stührwohldt et al., 2020 |
| pGGD011 | mGFP (GREEN FLUORESCENT PROTEIN; A206K) | Lupanga et al., 2020 |
| pGGD-GSL-mTurquoise | mTurquoise with GS linker | Waadt et al., 2017 |
| pGGD-GFP11 | ß-strand 11 of split GFP (GFP11) | This work |
| pGGD-GFP11x2 | 2-fold tandem repeat for ß-strand 11 of split GFP (GFP11x2) | This work |
| pGGE001 | RBCS terminator (pea) | Lampropoulos, et al., 2013 |
| pGGF Alli-mCherry | Seed coat marker cassette for mCherry | Stephani et al., 2020 |
| pGGF Alli-YFP | Seed coat marker cassette for YFP | Stephani et al., 2020 |
| pGGG004 | FH adapter for second expression cassette | Lampropoulos, et al., 2013 |
| pGGG005 | HA adapter for second expression cassette | Lampropoulos, et al., 2013 |

**Table S6. Primers for amplification of *amiR-vps16* sequence.**

| Primer name | Primer sequence |
|---|---|
| Primer A | CTGCAAGGCGATTAAGTTGGGTAAC |
| Primer B | GCGGATAACAATTTCACACAGGAAACAG |
| I miR-s (vps16) | GATAAGTACTCAGATATCCGCCGCTCTCTTTTGTATTCCA |
| II miR-a (vps16) | AGCGGCGGATATCTGAGTACTTATCAAAGAGAATCAATGA |
| III miR*s (vps16) | AGCGACGGATATCTGTGTACTTTTCACAGGTCGTGATATG |
| IV miR*a (vps16) | GAAAAGTACACAGATATCCGTCGCTACATATATATTCCTA |

**Table S7. Primers used to generate pHEE-mCherry.**

| Primer name | Primer sequence |
|---|---|
| EcoRI-pHEE401E_fw1 | CAACGAATTCGTAATCATGTCATAGCTG |
| BsaI-pHEE401E_rev1 | AACAGGTCTCTGGCAAGCTGCTCTAGCCAATAC |
| BsaI-SCSC_fw | AACAGGTCTCATGCCTTGAAACCAAATTAACATAGGG |
| BsaI-SCSC_rev | AACAGGTCTCTGGGGGGTACCCCTGGATTTTGG |
| BsaI-pHEE401E_fw2 | AACAGGTCTCACCCCGAATTAATTCGGCGTTAATTC |
| SacII-pHEE401E_rev2 | GAAACCGCGGTGATCACAGG |

**Table S8. Primers used for the generation of CRISPR/Cas9 constructs.** Bases of incorporated gRNAs are bold.

| Primer name | Primer sequence |
|---|---|
| gRNA_NET4B_3_FW | ATATATGGTCTCGATT**GGAGCAAGAAGCAGTTTAAG**GTTTTAGAGCTAGAAATAGC |
| gRNA_NET4B_4_REV | ATTATTGGTCTCGAAAC**CATGTATCGCGCATTGGCTC**AATCTCTTAGTCGACTCTAC |
| gRNA_NET4A_1_FW | ATATATGGTCTCGATT**GGTGTTTCCAGCTTTCCCAT**GTTTTAGAGCTAGAAATAGC |
| gRNA_NET4A_2_REV | ATTATTGGTCTCGAAAC**CGATTTCGCTCTTCAGCTTC**AATCTCTTAGTCGACTCTAC |
| gRNA_NET4A_3_FW | ATATATGGTCTCGATT**GGTACCGTGCTTTGGCAGAG**GTTTTAGAGCTAGAAATAGC |
| gRNA_NET4A_4_REV | ATTATTGGTCTCGAAAC**TTCTGCTTAACGTCTCTATC**AATCTCTTAGTCGACTCTAC |
| gRNA_NET3C_1_FW | ATATATGGTCTCGATT**GATCGCTCTCAGAATACAAT**GTTTTAGAGCTAGAAATAGC |
| gRNA_NET3C_2_REV | ATTATTGGTCTCGAAAC**TGTCTAAGAGCTGTTTCCTC**AATCTCTTAGTCGACTCTAC |
| gRNA_NET3C_3_FW | ATATATGGTCTCGATT**GGAGCTCCAAACATTCTCAA**GTTTTAGAGCTAGAAATAGC |
| gRNA_NET3C_4_REV | ATTATTGGTCTCGAAAC**CCATTCTCTCAAGCAACGCC**AATCTCTTAGTCGACTCTAC |
| gRNA_NET3A_1_FW | ATATATGGTCTCGATT**GGATGGACTCATCAAAATGG**GTTTTAGAGCTAGAAATAGC |
| gRNA_NET3A_2_REV | ATTATTGGTCTCGAAAC**CTCTGTCTCCTCTAAAGAGC**AATCTCTTAGTCGACTCTAC |
| gRNA_NET3A_3_FW | ATATATGGTCTCGATT**GAATTACCAACCTCGGTGCG**GTTTTAGAGCTAGAAATAGC |
| gRNA_NET3A_4_REV | ATTATTGGTCTCGAAAC**CCTTCCTTATTACGCCTTGC**AATCTCTTAGTCGACTCTAC |

**Table S9. List of used CRISPR/Cas9 constructs for generating deletion mutants.** The gRNAs are given without “NGG”. Red bases were changed to G (gRNA 1 or 3) or C (gRNA 2 or 4) when incorporated into the primers listed in Table S6.

| **Construct** | **gRNA 1 or 3** | **gRNA 2 or 4** | **Base vector** |
|---|---|---|---|
| pSK2 | AGAGCAAGAAGCAGTTTAAG | CAGCCAATGCGCGATACATG | pHEE401E |
| pSK5 | TGTGTTTCCAGCTTTCCCAT | AAAGCTGAAGAGCGAAATCG | pHEE401E |
| pSK6 | TGTACCGTGCTTTGGCAGAG | CATAGAGACGTTAAGCAGAA | pHEE401E |
| pSK17 | TATCGCTCTCAGAATACAAT | GAGGAAACAGCTCTTAGACA | pHEE-mCherry |
| pSK18 | AGAGCTCCAAACATTCTCAA | TGCGTTGCTTGAGAGAATGG | pHEE-mCherry |
| pSK9 | TGATGGACTCATCAAAATGG | ACTCTTTAGAGGAGACAGAG | pHEE401E |
| pSK10 | AAATTACCAACCTCGGTGCG | GCAAGGCGTAATAAGGAAGG | pHEE401E |

**Table S10. List of CRISPR/Cas9-derived deletion mutants.** Base lines and constructs used to obtain the specific gene deletions are given for each line. All mutant lines are homozygous for the introduced deletion and lost the integrated CRISPR/Cas9 construct during the screening process.

| **Deletion mutant** | **Base line** | **Construct** |
|---|---|---|
| *net4b* (CRISPR mutant) | Col-0 | pSK2 |
| *rabg3a,b,c,d,e,f net4a #1* | *rabg3a,b,c,d,e,f* (Ebine et al., 2014) | pSK6 |
| *rabg3a,b,c,d,e,f net4a* #2 | *rabg3a,b,c,d,e,f* (Ebine et al., 2014) | pSK5 |
| *net3c* | Col-0 | pSK18 |
| *net3c net4a net4b #2* | *net4b* (CRISPR mutant, this work) | pSK17 & pSK5 |
| *net3a* | Col-0 | pSK9 |
| *net3a net4a net4b #1* | *net4a net4b* (T-DNA mutant, Kaiser et al., 2019) | pSK10 |
| *net3a net4a net4b #2* | *net4a net4b* (T-DNA mutant, Kaiser et al., 2019) | pSK9 |

**Table S11. Primers used for the generation of rBiFC and mbSUS constructs.**

| Primer name | Primer sequence | Purpose |
|---|---|---|
| attB1_NET4A.FOR | GGGGACAAGTTTGTACAAAAAAGCAGGCTCTATGGATTATGATCTGCTTCGTTCCAAGA | Amplification of sequences for rBiFC constructs (used entry vector: pDONR221-P1P4, used destination vector: pBiFCt-2in1-NC) |
| attB4-wo_NET4A.REV | GGGGACAACTTTGTATAGAAAAGTTGGGTGAGAAGCAAGAATGGATGATGGTCTTG | |
| attB3_RABG3b.FOR | GGGGACAACTTTGTATAATAAAGTTGCTATGTCGACGCGAAGACGAAC | Amplification of sequences for rBiFC constructs (used entry vector: pDONR221-P3P2, used destination vector: pBiFCt-2in1-NC) |
| attB2-ST_RABG3b.REV | GGGGACCACTTTGTACAAGAAAGCTGGGTTCAGCAAGCACAACCTCCTCT | |
| attB3_RABG3a.FOR | GGGGACAACTTTGTATAATAAAGTTGCTATGGCGACGAGAAGACGTAC | |
| attB2-ST_RABG3a.REV | GGGGACCACTTTGTACAAGAAAGCTGGGTTCAGCAAGCGCAACCACC | |
| attB3_VAMP711.FOR | GGGGACAACTTTGTATAATAAAGTTGCTATGGCGATTCTGTACGCCC | |
| attB2-ST_VAMP711.REV | GGGGACCACTTTGTACAAGAAAGCTGGGTTTAAATGCAAGATGGTAGAGTAGGTCCG | |
| attB3_RABC1.FOR | GGGGACAACTTTGTATAATAAAGTTGCTATGGGTTCTTCGTCAGGACAAC | |
| attB2-ST_RABC1.REV | GGGGACCACTTTGTACAAGAAAGCTGGGTCTAAGACGAGCAGCAGTAGCT | |
| attB3_1245.FOR | GGGGACAACTTTGTATAATAAAGTTGCTATGGTGATTCACCTTAGTTCATCTTGTTATTCC | |
| attB2-ST_1245.REV | GGGGACCACTTTGTACAAGAAAGCTGGGTTCATGATGTACAACAAAAGTACCATTGCTG | |
| attB1_RABG3b.FOR | GGGGACAAGTTTGTACAAAAAAGCAGGCTCTATGTCGACGCGAAGACGAAC | Amplification of sequences for mbSUS prey constructs (used entry vector: pDONR207, used destination vector: pNX35-DEST-1) |
| attB2-ST_RABG3b.REV | GGGGACCACTTTGTACAAGAAAGCTGGGTTCAGCAAGCACAACCTCCTC | |
| attB1_RABG3a.FOR | GGGGACAAGTTTGTACAAAAAAGCAGGCTCTATGGCGACGAGAAGACGTAC | |
| attB2-ST_RABG3a.REV | GGGGACCACTTTGTACAAGAAAGCTGGGTTCAGCAAGCGCAACCACCG | |
| attB1_RABC1.FOR | GGGGACAAGTTTGTACAAAAAAGCAGGCTCTATGGGTTCTTCGTCAGGAC | |
| attB2-ST_RABC1.REV | GGGGACCACTTTGTACAAGAAAGCTGGGTCTAAGACGAGCAGCAGTAGCTC | |
| attB1_1245.FOR | GGGGACAAGTTTGTACAAAAAAGCAGGCTCTATGGTGATTCACCTTAGTTCATCTTG | |
| attB2-ST_1245.REV | GGGGACCACTTTGTACAAGAAAGCTGGGTTCATGATGTACAACAAAAGTACCATTGC | |
| attB1_15042.FOR | GGGGACAAGTTTGTACAAAAAAGCAGGCTCTATGGACGCTACCAACTTTGGAC | |
| attB2-ST_15042.REV | GGGGACCACTTTGTACAAGAAAGCTGGGTTTAACGAGTTGTGCTGCTTGTG | |
| attB1_VAMP711.FOR | GGGGACAAGTTTGTACAAAAAAGCAGGCTTAATGGCGATTCTGTACGCCCT | |
| attB2-ST_VAMP711.REV | GGGGACCACTTTGTACAAGAAAGCTGGGTTTAAATGCAAGATGGTAGAGTAGG | |
| attB1_VAP27-1.FOR | GGGGACAAGTTTGTACAAAAAAGCAGGCTCTATGAGTAACATCGATCTGATTGGGATGAGTAAC | |
| attB2-ST_VAP27-1.REV | GGGGACCACTTTGTACAAGAAAGCTGGGTTTATGTCCTCTTCATAATGTATCCCAAAATTAGACC | |
| attB1_ORP1C.FOR | GGGGACAAGTTTGTACAAAAAAGCAGGCTCTATGCATCCCTTCTGTTGCGT | |
| attB2-ST_ORP1C.REV | GGGGACCACTTTGTACAAGAAAGCTGGGTTTACTCTATCATTTGTTGATCGGAATCGATGT | |

| | | |
|---|---|---|
| attB1_ORP2A.FOR | GGGGACAAGTTTGTACAAAAAAGCAGGCTCTATGCGGGTTAAAGAGTTACATCCAC | |
| attB2-ST_ORP2A.REV | GGGGACCACTTTGTACAAGAAAGCTGGGTCTAAGCGGCGTCCGCTAG | |
| attB1_ORP3C.FOR | GGGGACAAGTTTGTACAAAAAAGCAGGCTCTATGGGAAGTCCAAAGAAGAACGAGA | |
| attB2-ST_ORP3C.REV | GGGGACCACTTTGTACAAGAAAGCTGGGTTCATTCGGTTGAGATATTACCATATTGCCAAG | |
| attB1_NET4A.FOR | GGGGACAAGTTTGTACAAAAAAGCAGGCTCTATGGATTATGATCTGCTTCGTTCCAAGA | Amplification of sequences for mbSUS bait constructs (used entry vector: pDONR207, used destination vector: pMETOYC-Dest) |
| attB2-wo_NET4A.REV | GGGGACCACTTTGTACAAGAAAGCTGGGTGAGAAGCAAGAATGGATGATGGTC | |
| attB1_NET4B.FOR | GGGGACAAGTTTGTACAAAAAAGCAGGCTCTATGGCTTCGTCTACGGCTCAG | |
| attB2-wo_NET4B.REV | GGGGACCACTTTGTACAAGAAAGCTGGGTGAGTTGATAAGACCACTACTCTCTTACTCTTATGT | |
| attB1_NET3A.FOR | GGGGACAAGTTTGTACAAAAAAGCAGGCTCTATGGTGATGGACTCATCAAAATGGTG | |
| attB2-wo_NET3A.REV | GGGGACCACTTTGTACAAGAAAGCTGGGTGAAGAGTCATGAGCTCTTTGGAAGTAG | |
| attB1_NET3C.FOR | GGGGACAAGTTTGTACAAAAAAGCAGGCTCTATGGTTAGAGAAGAGGAGAAATCGAGATG | |
| attB2-wo_NET3C.REV | GGGGACCACTTTGTACAAGAAAGCTGGGTGAAGGACCTTGTTGCCATCGC | |

**Table S12: List of generated Gateway expression constructs used for rBiFC and mbSUS experiments.** The used destination vectors, including tags, as well as the sequences cloned from suitable entry vectors are shown for each construct.

| Constructs | N-terminal tag | Coding (or genomic) sequence | C-terminal tag | Destination vector |
|---|---|---|---|---|
| NET4A-cYFP+nYFP-VAMP711 | nYFP-HA | NET4A + RFP + VAMP711 | MYC-cYFP | pBiFCt-2in1-NC |
| NET4A-cYFP+nYFP-RABG3b | nYFP-HA | NET4A + RFP + RABG3b | MYC-cYFP | pBiFCt-2in1-NC |
| NET4A-cYFP+nYFP-RABG3a | nYFP-HA | NET4A + RFP + RABG3a | MYC-cYFP | pBiFCt-2in1-NC |
| NET4A-cYFP+nYFP-RABC1 | nYFP-HA | NET4A + RFP + RABC1 | MYC-cYFP | pBiFCt-2in1-NC |
| NET4A-cYFP+nYFP-gAt4g01245 | nYFP-HA | NET4A + RFP + At4g01245 (genomic) | MYC-cYFP | pBiFCt-2in1-NC |
| NubG-RABG3b | NubG-2xHA | RABG3b | - | pNX35-DEST-1 |
| NubG-RABG3a | NubG-2xHA | RABG3a | - | pNX35-DEST-1 |
| NubG-RABC1 | NubG-2xHA | RABC1 | - | pNX35-DEST-1 |
| NubG-At2g15042 | NubG-2xHA | At2g15042 | - | pNX35-DEST-1 |
| NubG-gAt4g01245 | NubG-2xHA | At4g01245 (genomic) | - | pNX35-DEST-1 |
| NubG-VAMP711 | NubG-2xHA | VAMP711 | - | pNX35-DEST-1 |
| NET4A-Cub | Ost4 | NET4A | Cub | pMETOYC-Dest |
| NET4B-Cub | Ost4 | NET4B | Cub | pMETOYC-Dest |
| NET3A-Cub | Ost4 | NET3A | Cub | pMETOYC-Dest |
| NET3C-Cub | Ost4 | NET3C | Cub | pMETOYC-Dest |
| NubG-VAP27-1 | NubG-2xHA | VAP27-1 | - | pNX35-DEST-1 |
| NubG-ORP1C | NubG-2xHA | ORP1C | - | pNX35-DEST-1 |
| NubG-ORP2A | NubG-2xHA | ORP2A | - | pNX35-DEST-1 |
| NubG-ORP3C | NubG-2xHA | ORP3C | - | pNX35-DEST-1 |

**Table S13. Primers used to clone constructs for recombinant protein expression.**

| Primer name | Primer sequence |
|---|---|
| NET4A_fw_BtgZI | AACAGCGATGTCTAGGACGACATGGATTATGATCTGCTTCGTTCCAAGAAG |
| NET4A1/2_rev_XhoI | ATCTCTCGAGAGAAGCAAGAATGGATGATGGTCTTGTTG |

**Table S14. List of Arabidopsis lines obtained by crossing.** The parental lines used for crossing and information about the homozygosity are given for each line.

| **Crossing line** | **Mother line** | **Father line** | **Status** |
|---|---|---|---|
| lifeact-RFP x NET4A-GFP$^{OE}$ | lifeact-RFP | NET4A-GFP$^{OE}$ | homozygous |
| *act7-4* x NET4A-GFP$^{OE}$ | *act7-4* | NET4A-GFP$^{OE}$ | homozygous |
| *act2 act8* x NET4A-GFP$^{OE}$ | *act2 act8* | NET4A-GFP$^{OE}$ | homozygous |
| NET4A-GFP$^{OE}$ x *amiR-vps16* | NET4A-GFP$^{OE}$ | *amiR-vps16* | homozygous |
| SRß-mTurquoise x NET4A-GFP$^{OE}$ | SRß-mTurquoise | NET4A-GFP$^{OE}$ | heterozygous |
| RFP-HDEL x NET4A-GFP | RFP-HDEL | NET4A-GFP | heterozygous |
| RFP-HDEL x NET4A-GFP$^{OE}$ | RFP-HDEL | NET4A-GFP$^{OE}$ | heterozygous |

**Table S15. Primers used for genotyping.** For T-DNA insertional mutants, primers were used as described previously in Kandasamy et al., 2009 or designed and used as recommended by the Nottingham Arabidopsis Stock Centre (NASC) (Nottingham University, UK; http://signal.salk.edu/tdnaprimers.2.html).

| **Primer name** | **Primer sequence** | **Purpose** |
|---|---|---|
| GT_amiR-vps16_fw | CACGCTCGGACGCATATTAC | Detection of integrated *amiR-vps16* construct |
| GT_amiR-vps16_rev | GCGGCGGATATCTGAGTACTTATC | |
| GFP_fw | CCTCGTGACCACCTTCACCTAC | Detection of integrated GFP-fusion constructs |
| GFP_rev | GTGATCGCGCTTCTCGTTGG | |
| GT_NET4B-A_fw | GGCCGCGTTTCCAATTATC | Screening for CRISPR/Cas9-provoked gene deletions |
| GT_NET4B-B_rev | GCTAGATTGACGGCGACTC | |
| GT_NET4A_fw | CACTTGGCCACGTTGAAATTGC | |
| GT_NET4A_rev | GGCTATGCGTAGTCTCTTGTATTCG | |
| GT_NET3C_fw | GGTTGAAGACGAGCGACATGG | |
| GT_NET3C_rev | GCTAGTGCGCAGTTGTTGTACTTG | |
| GT_NET3A_fw | CTGAACTCTCAAGTCCGAAGACAG | |
| GT_NET3A_rev | CAATCCTATGATATACCTTACGCTAAC | |
| EC_fw | CGTCTCCAATAGGAGCGCTACTG | Screening for integrated CRISPR/Cas9 constructs |
| EC_rev | GGGCCGATTAGAATCACTCAGTC | |
| ACT2-173S | CTTCCTCAATCTCATCTTCTT | Screening for T-DNA insertions |
| ACT2-AS | CATGACACCATGATGTCTTGGCCT | |
| ACT2-173S_LBp | GCTCAGGATCCGATTGTCGTTTCCCGCCTT | |
| ACT8-5'S4 | TCGATCAAGATTCAGATCTTTATG | |
| ACT8-74A | ACACCATGCTCAATAGGGTATTTCAAT | |
| LB-GABIS1 | CCCATTTGGACGTGAATGTAGACAC | |
| AAc7S | AGGATTCTTCTCGCTTCTGTCGATCTCTCGCT | |
| AAc7-I1A | AAATCATGATCAGTAGTCTTACACATGT | |
| JLB7804 | TTGGTAATTACTCTTTCTTTTCCTCCATATT | |
