## Supplemental figures for "NETWORKED proteins link actin-dependent vacuolar organization to the endoplasmic reticulum"

### Supplementary Figures

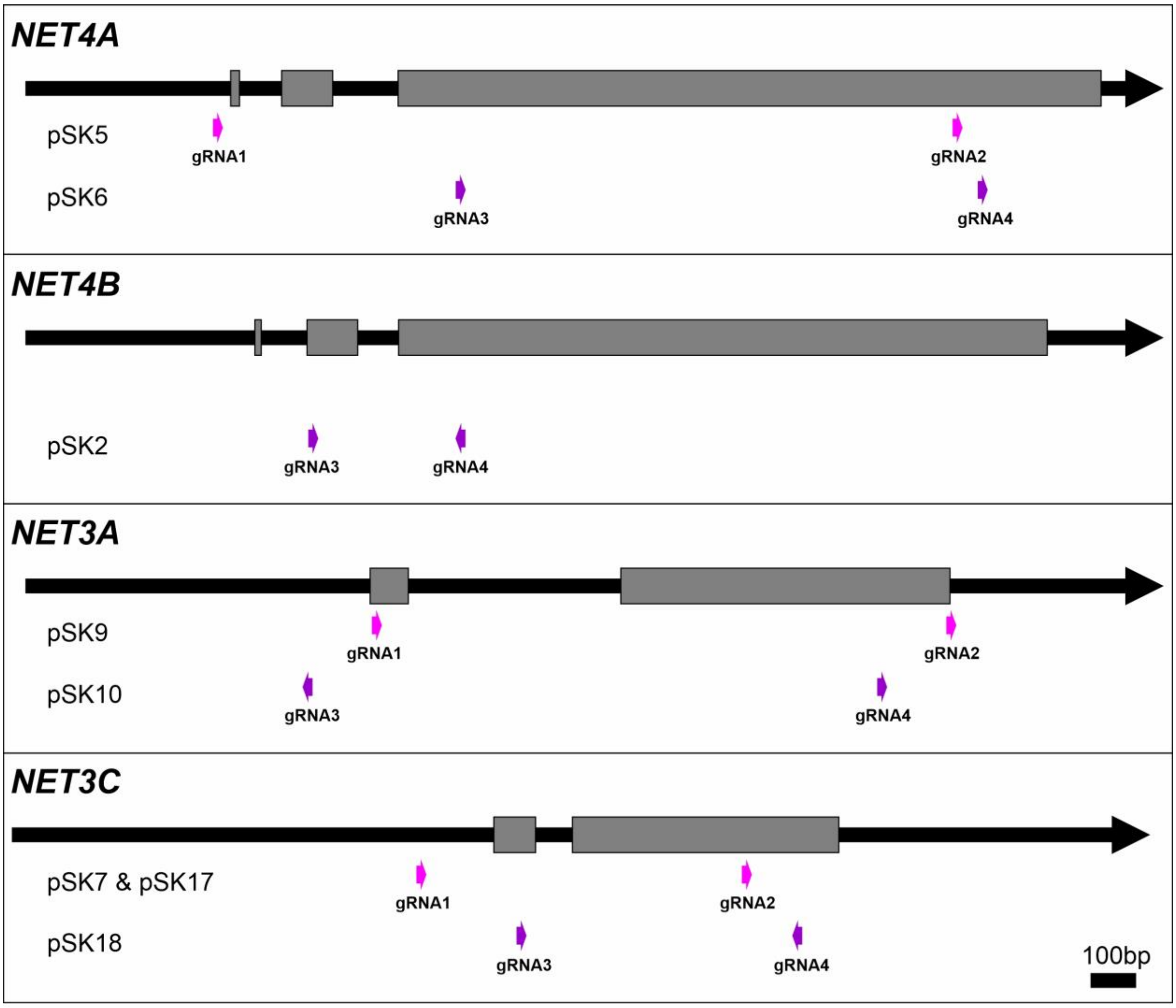


| Mutant | gRNAs / source |
|---|---|
| *net4b* | gRNA3 + gRNA4 |
| *net3a* | gRNA1 + gRNA2 |
| *net3c* | gRNA3 + gRNA4 |
| *net3a net4a net4b* #1 | *NET3A*: gRNA3 + gRNA4 / *net4a net4b* from Kaiser et al. (2019) |
| *net3a net4a net4b* #2 | *NET3A*: gRNA1 + gRNA2 / *net4a net4b* from Kaiser et al. (2019) |
| *net3c net4a net4b* | *NET3C*: gRNA1 + gRNA2; *NET4A*: gRNA1 + gRNA2 / *net4b* from this study |
| *net4a net4b* | Kaiser et al. (2019) |
| *rabg3a,b,c,d,e,f net4a* #1 | *NET4A*: gRNA3 + gRNA4 |
| *rabg3a,b,c,d,e,f net4a* #2 | *NET4A*: gRNA1 + gRNA2 |

**Fig. S1. Schematic overview of CRISPR/Cas9-derived deletions for *NET4A*, *NET4B*, *NET3A* and *NET3C*.** The genomic region of each gene is illustrated as black arrow showing exons as grey bars. The genomic sites of each gRNA pair used to generate CRISPR/Cas9-based deletion mutants are shown as coloured arrows. For each gene, gRNA1 and 2 (pink arrows) or gRNA3 and 4 (violet arrows) were used in combination to delete the genomic area in between the two gRNAs. The generated plasmids, which contain the individual gRNA pairs are given on the left site and details listed in Table S9. Details on all deletion mutants established using the presented plasmids/gRNA pairs are given in Table S10. Arrow heads of gRNAs mark their direction.

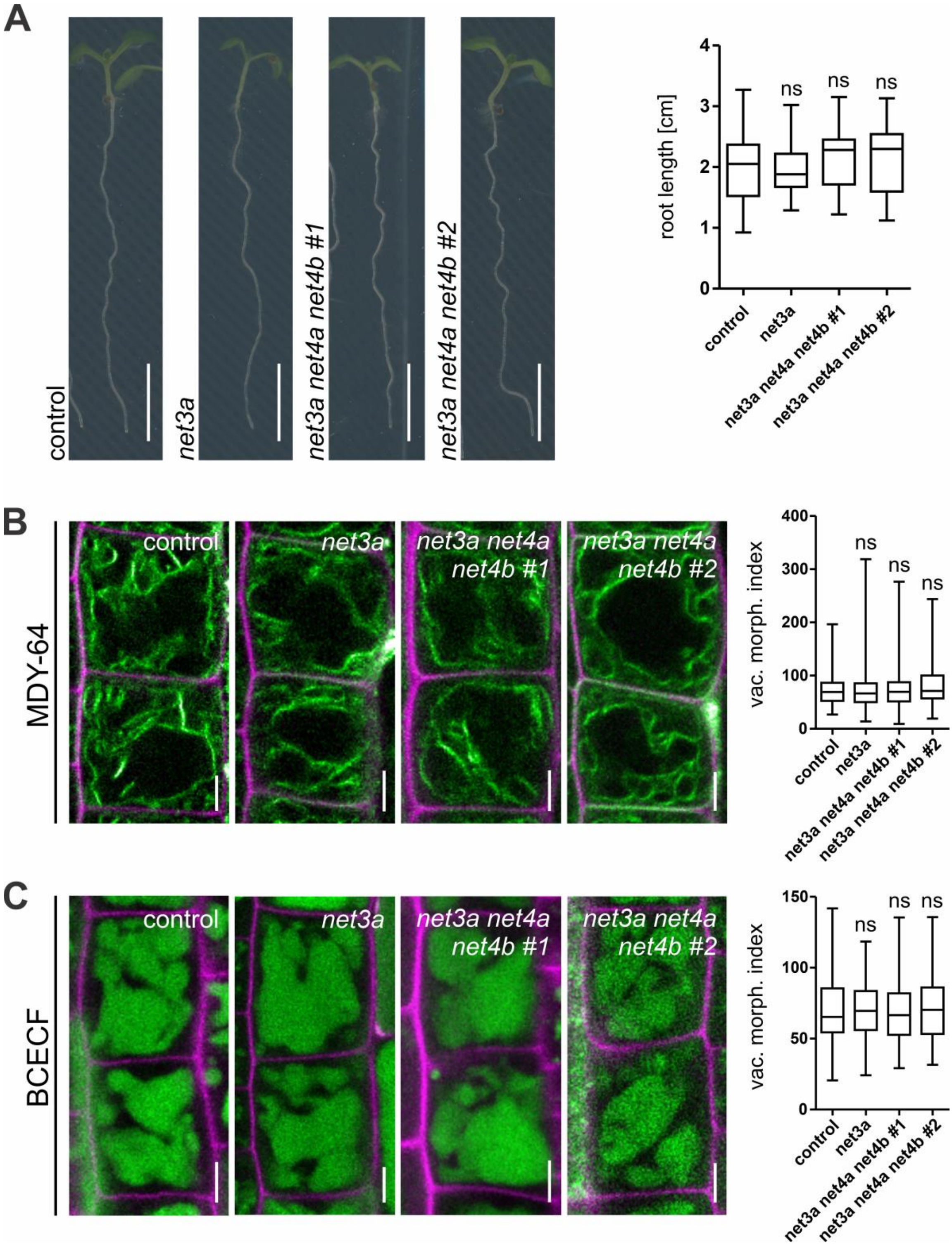


**Fig. S2. Phenotypic characterization of *net3a* and *net4a net4b net3a* mutants. (A)** Investigation of root length for lines with CRISPR/Cas9 derived deletion of *NET3A* in Col-0 wild type (n=55) and *net4a net4b* t-DNA mutant (*net3a net4a net4b #1* (n=56) and *net3a net4a net4b #2* (n=61)) compared to Col-0 control (n=150). Scale bars = 0.5 cm. **(B)** Analysis of vacuolar morphology based on the tonoplast dye MDY-64 (n=88 for control, n=100 for *net3a*, n=108 for *net3a net4a net4b #1* and n=88 for *net3a net4a net4b #2*). **(C)** Analysis of vacuolar morphology based on the vacuole dye BCECF (n=96 for control and *net3a*, n=100 for *net3a net4a net4b #1* and n=108 for *net3a net4a net4b #2*). Box limits represent 25th-75th percentile, the horizontal line the median and whiskers minimum to maximum values. Differences were analyzed by one-way ANOVA and Dunnett's post hoc test; ns = not significant. Scale bars = 5 µm.

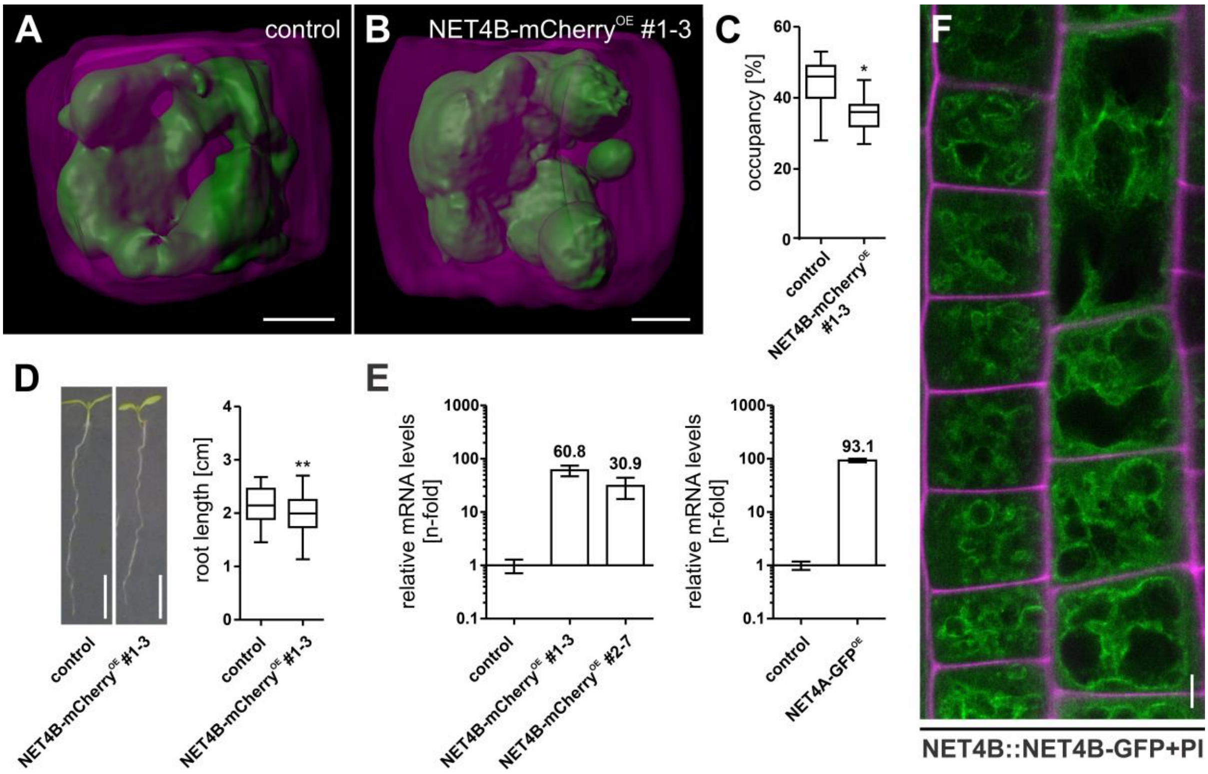


**Fig. S3. NET4B functions similar to NET4A. (A)** 3D vacuole models of Col-0 wild type control and **(B)** NET4B-mCherry$^{OE}$ epidermal cells from the root meristem. **(C)** Quantification of vacuolar occupancy of the cell (n=7 for each line). **(D)** Root length of NET4B-mCherry$^{OE}$ (n=62) in comparison to the control (n=50). Significant differences were analyzed by Student's t-test, *$P < 0.05$; **$P < 0.01$. **(E)** Investigation of gene upregulation in NET4B and NET4A overexpression lines by qRT-PCR. **(F)** Subcellular localization of NET4B-GFP (green), expressed under the endogenous promoter. Cell walls (magenta) were stained with propidium iodide (PI). Scale bars = 5 µm (A, B, F) or 5 mm (D).

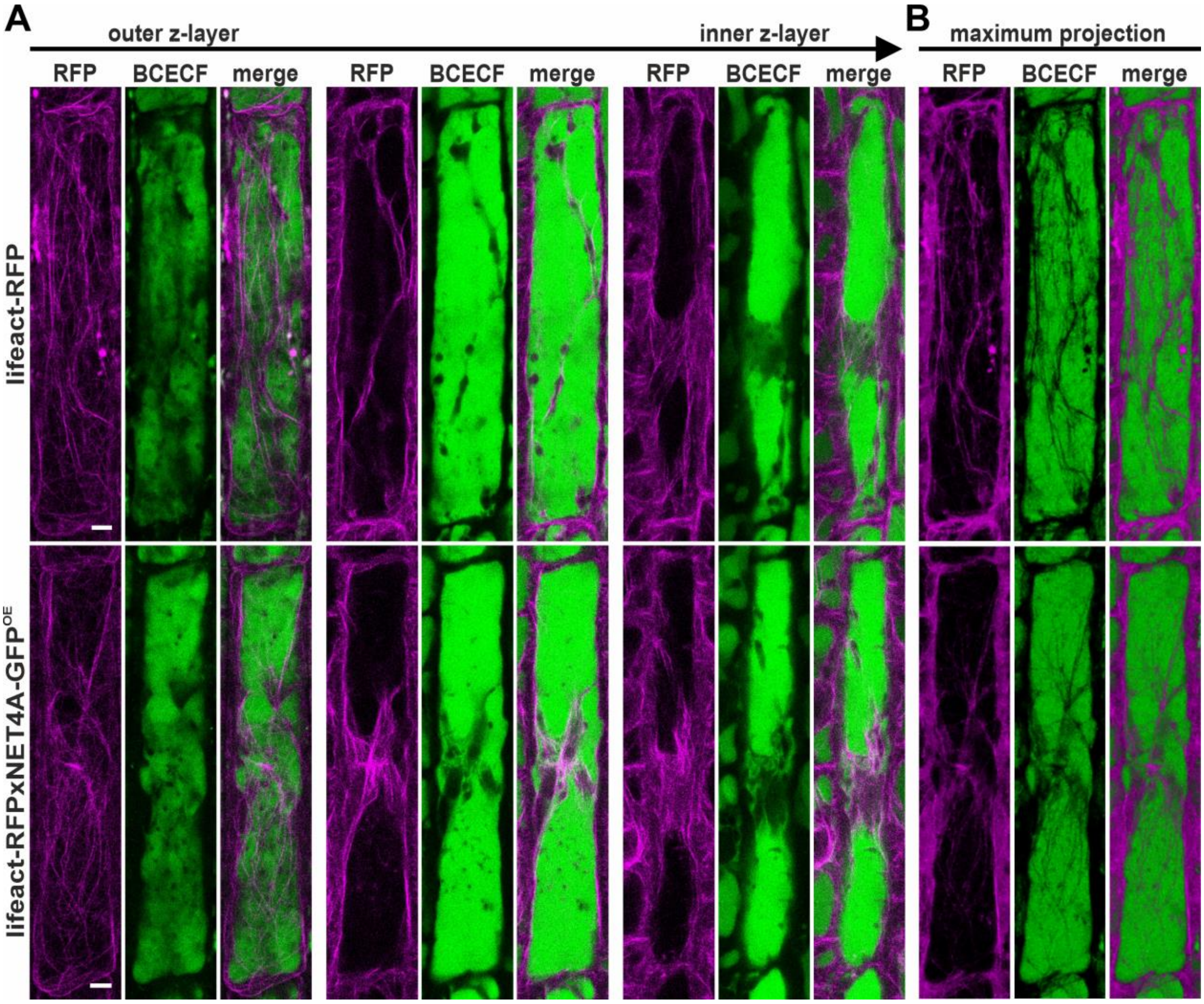


**Fig. S4**. **NET4A overexpression leads to actin bundling from the cell periphery to the nucleus. (A)** Actin filaments decorated by lifeact-RFP (magenta) and BCECF-stained vacuoles (green) of lifeact-RFP and lifeact-RFP x NET4A-GFP$^{OE}$ for different optical sections starting from the cortex towards the centre of the cell. **(B)** Maximum projections of (A). Scale bars = 5 µm.

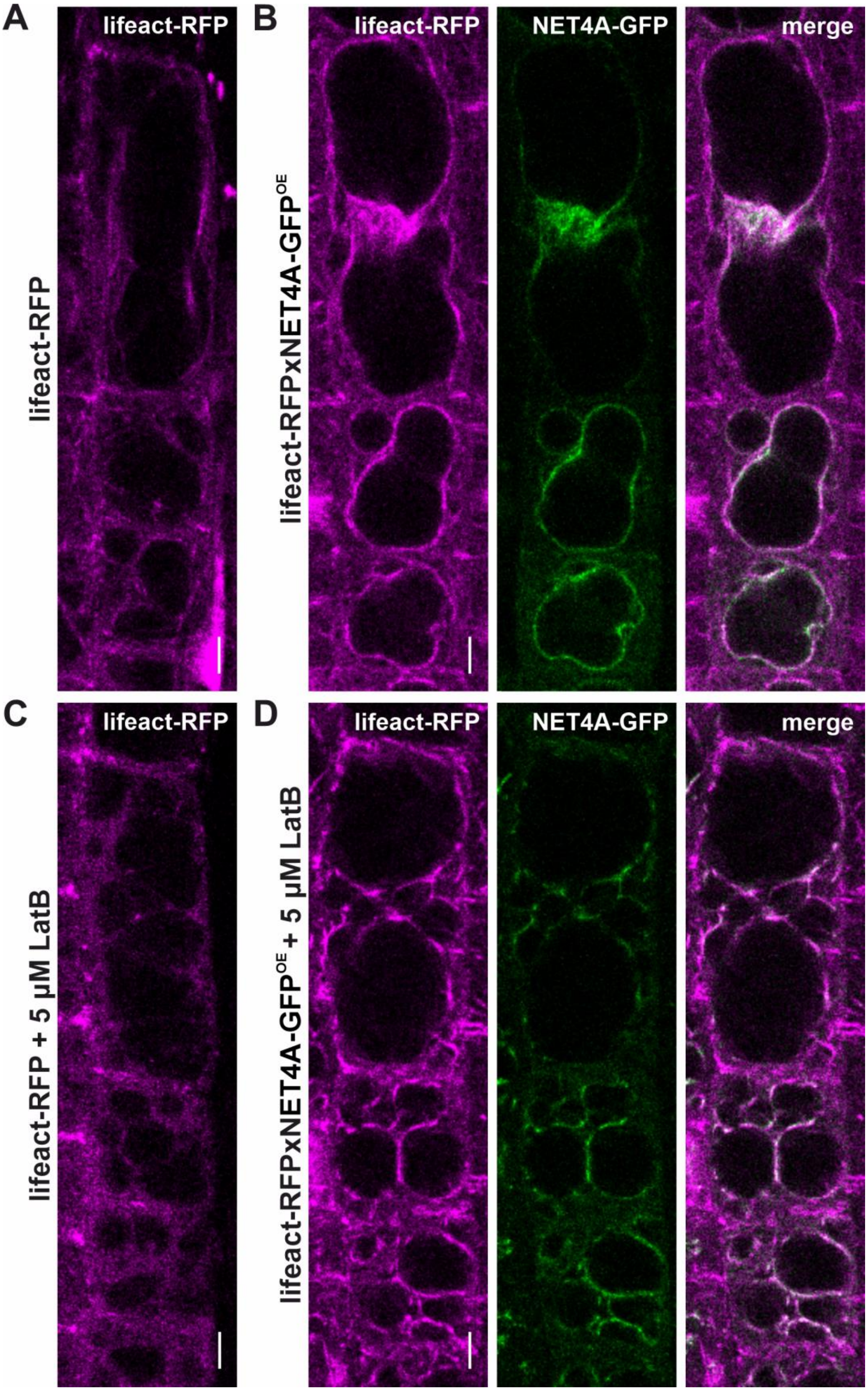


**Fig. S5: NET4A overexpression stabilizes actin filaments. (A)** lifeact-RFP (magenta) signals in epidermal cells of the root transition zone. **(B)** Transversal optical sections of cells expressing lifeact-RFP and NET4A-GFP$^{OE}$ (green). **(C)** LatB treatment (5 µM, 5h) of lifeact-RFP. **(D)** Same treatment as in (C) of lifeact-RFP x NET4A-GFP$^{OE}$. Scale bars = 5 µm.

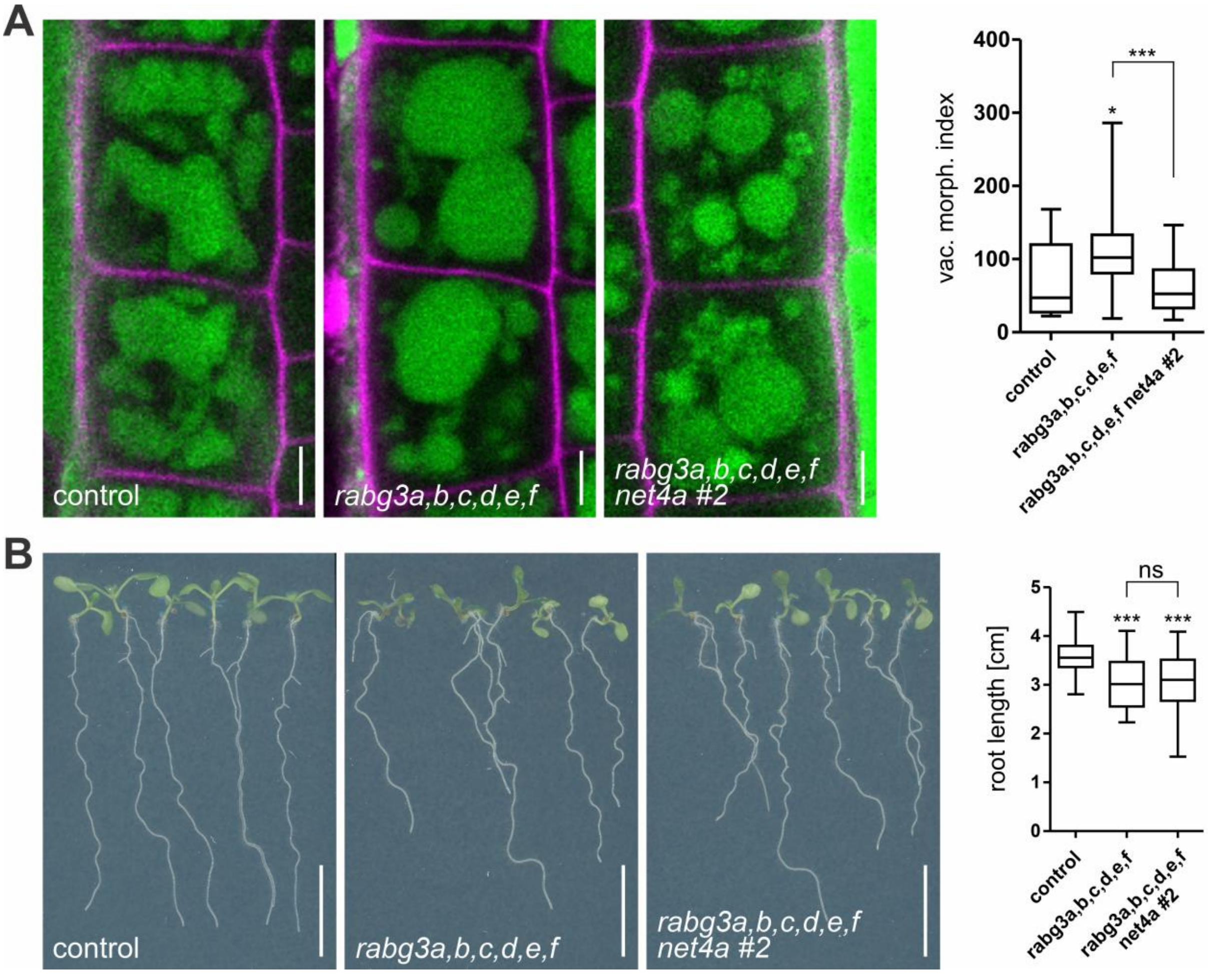


**Fig. S6.** ***rabg3a,b,c,d,e,f*** **sextuple and** ***rabg3a,b,c,d,e,f net4a*** **septuple mutants show more round and seemingly disconnected vacuoles and inhibited root growth. (A)** Vacuoles in the *rabg3a,b,c,d,e,f* sextuple (n=28) and the *rabg3a,b,c,d,e,f net4a* #2 septuple mutant (n=32) compared to Col-0 wild type control (n=28) and quantification of vacuolar morphology index. **(B)** 6 days old Arabidopsis seedlings. Root length of *rabg3a,b,c,d,e,f* (n=35) and *rabg3a,b,c,d,e,f net4a #2* (n=41) was quantified in comparison to the Col-0 control (n=48). Box limits represent 25th-75th percentile, the horizontal line the median and whiskers minimum to maximum values. Significant differences are shown (one-way ANOVA and Tukey post hoc test; ns = not significant; *P < 0.05; ****P* < 0.001). Scale bars = 5 µm for (A) and 1 cm for (B).

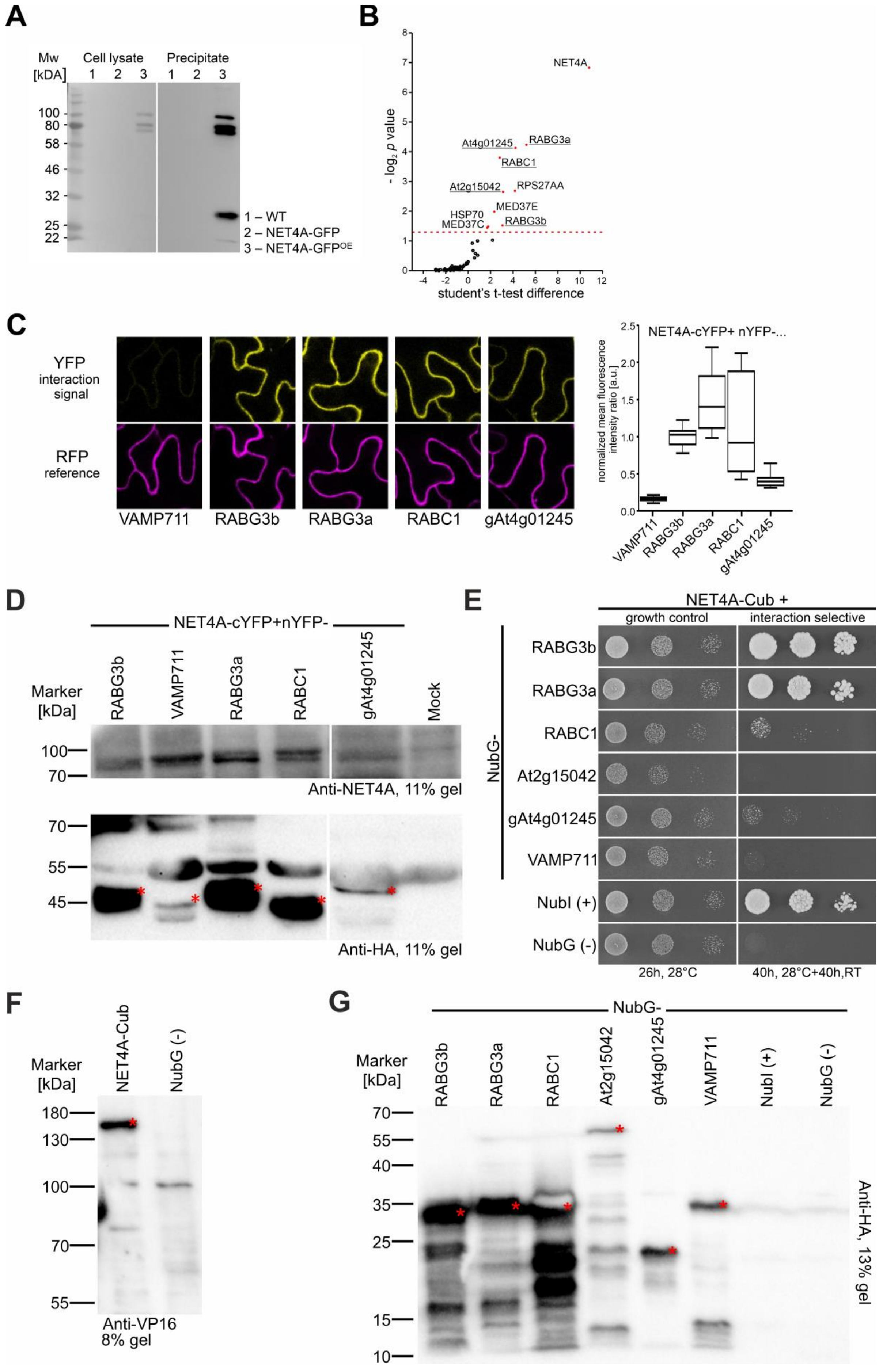

A
Mw [kDA]
Cell lysate
Precipitate
1 2 3 1 2 3
100
80
58
46
32
25
22
1 – WT
2 – NET4A-GFP
3 – NET4A-GFP^OE
B
NET4A
At4g01245
RABG3a
RABC1
At2g15042
RPS27AA
MED37E
HSP70
MED37C
RABG3b
- log2 p value
student's t-test difference
C
YFP interaction signal
RFP reference
VAMP711
RABG3b
RABG3a
RABC1
gAt4g01245
NET4A-cYFP+ nYFP-...
normalized mean fluorescence intensity ratio [a.u.]
D
NET4A-cYFP+nYFP-
Marker [kDa]
RABG3b
VAMP711
RABG3a
RABC1
gAt4g01245
Mock
Anti-NET4A, 11% gel
Anti-HA, 11% gel
E
NET4A-Cub +
growth control
interaction selective
NubG-
RABG3b
RABG3a
RABC1
At2g15042
gAt4g01245
VAMP711
NubI (+)
NubG (-)
26h, 28°C
40h, 28°C+40h,RT
F
Marker [kDa]
NET4A-Cub
NubG (-)
Anti-VP16
8% gel
G
NubG-
Marker [kDa]
RABG3b
RABG3a
RABC1
At2g15042
gAt4g01245
VAMP711
NubI (+)
NubG (-)
Anti-HA, 13% gel

**Fig. S7. Identification and confirmation of NET4A interaction partners. (A)** Detection of NET4A-GFP fusion protein for the indicated lines using a GFP specific antibody after co-immunoprecipitation of NET4A-GFP interaction partners with GFP-trap magnetic agarose beads. **(B)** LC-MS/MS-based identification of proteins that were co-immunoprecipitated with NET4A-GFP. The dashed red line marks the threshold for significant enrichment with NET4A-GFP compared to the wild type (control). Proteins used for confirmation are underlined. **(C)** Representative images and quantification of ratiometric bimolecular fluorescence complementation (rBiFC) measurements in *N. benthamiana* leaf epidermis cells. For quantification, YFP signals were normalized to the respective RFP signals to allow comparison of interaction strength. VAMP711, a SNARE localized to the tonoplast was used as negative control. **(D)** Confirmation of NET4A-cYFP (bait) and interaction partner (nYFP, prey) expression for rBiFC via immunological detection. **(E)** Mating-based Split-Ubiquitin System (mbSUS) growth assays in yeast using NET4A-Cub to test for interaction with NubG-RABG3a, NubG-RABG3b, NubG-RABC1, NubG-At2g15042 and NubG-gAt4g01245. For mbSUS growth assays THY.AP4 (MATa) yeast strains transformed with Cub-fusions were mated with THY.AP5 (MATα) strains transformed with NubG-fusions. Growth was assayed in dilution series of $OD_{600}$ from 1 to 0.001 for the given time and temperature on growth control media to confirm presence of vector fusion constructs and interaction selective media to test for specific interaction dependent activation of the reporter genes. NubG was used as negative, NubI as positive control. **(F)** Expression control of NET4A by immunological detection. **(G)** Western blot indicating NubG-construct expression. Expected size is highlighted by red asterisks.

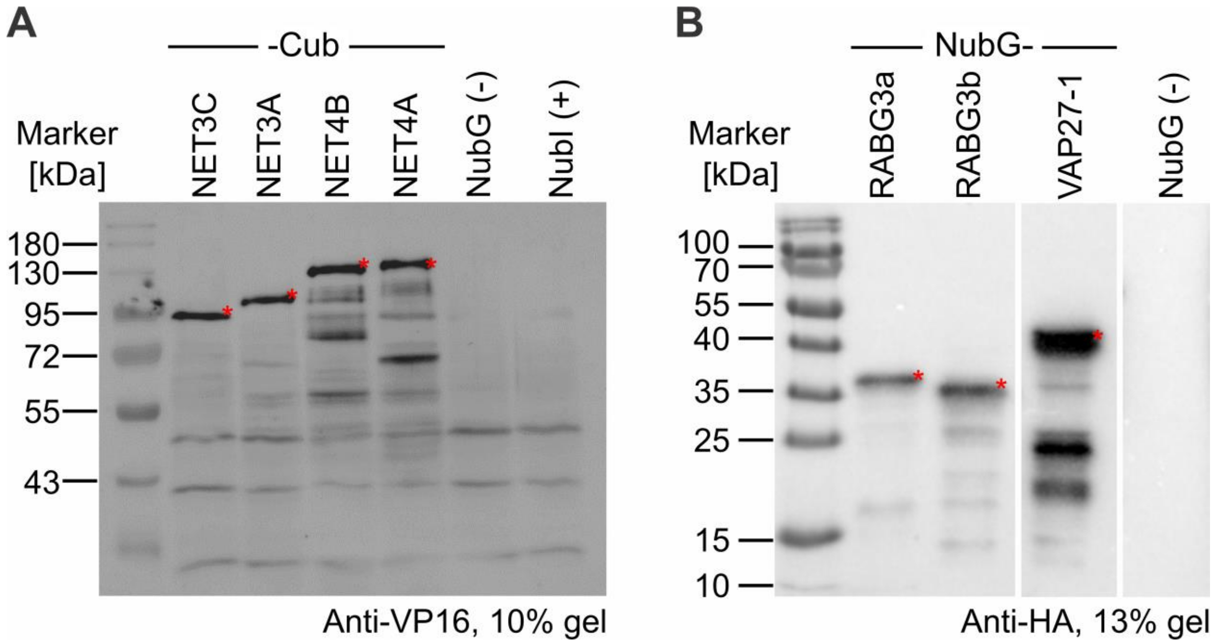


**Fig. S8. Expression control for mbSUS experiments in Fig. 6 and Fig. 7. (A)** Western blot showing detection of NET4A-, NET4B-, NET3A- and NET3C-Cub. **(B)** Western blot showing detection of NubG-RABG3a, -RABG3b and -VAP27-1. Expected size is highlighted by red asterisks.

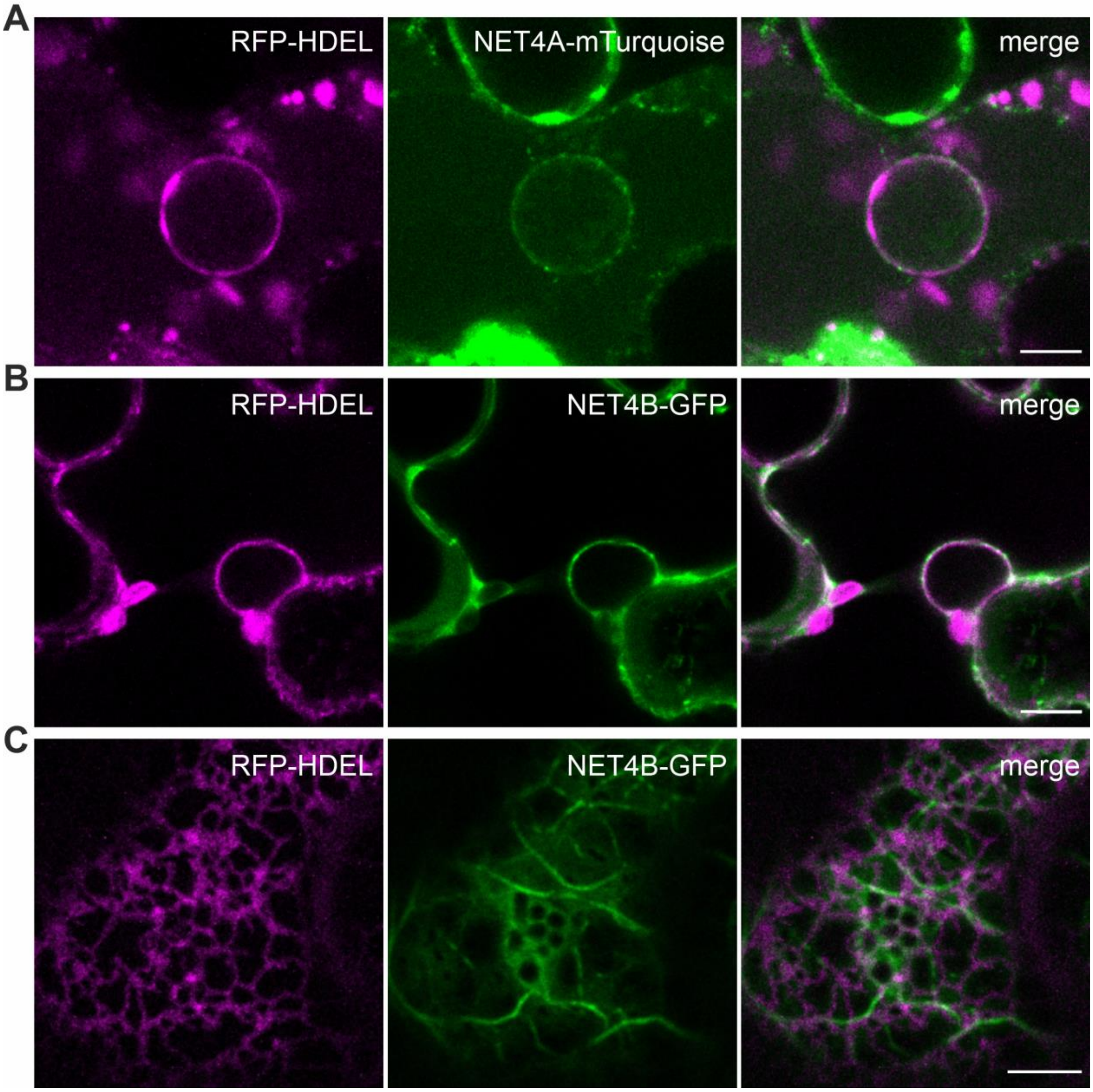


**Fig. S9. NET4A and NET4B both show localization at the nuclear envelope. (A)** and **(B)** Colocalization of NET4A-mTurquoise (A) or NET4B-GFP (B) with the ER marker RFP-HDEL at the nuclear envelope upon coexpression in epidermal cells of *N. benthamiana* leaves. **(C)** Cortical view of NET4B-GFP and RFP-HDEL. Scale bars = 5 μm.

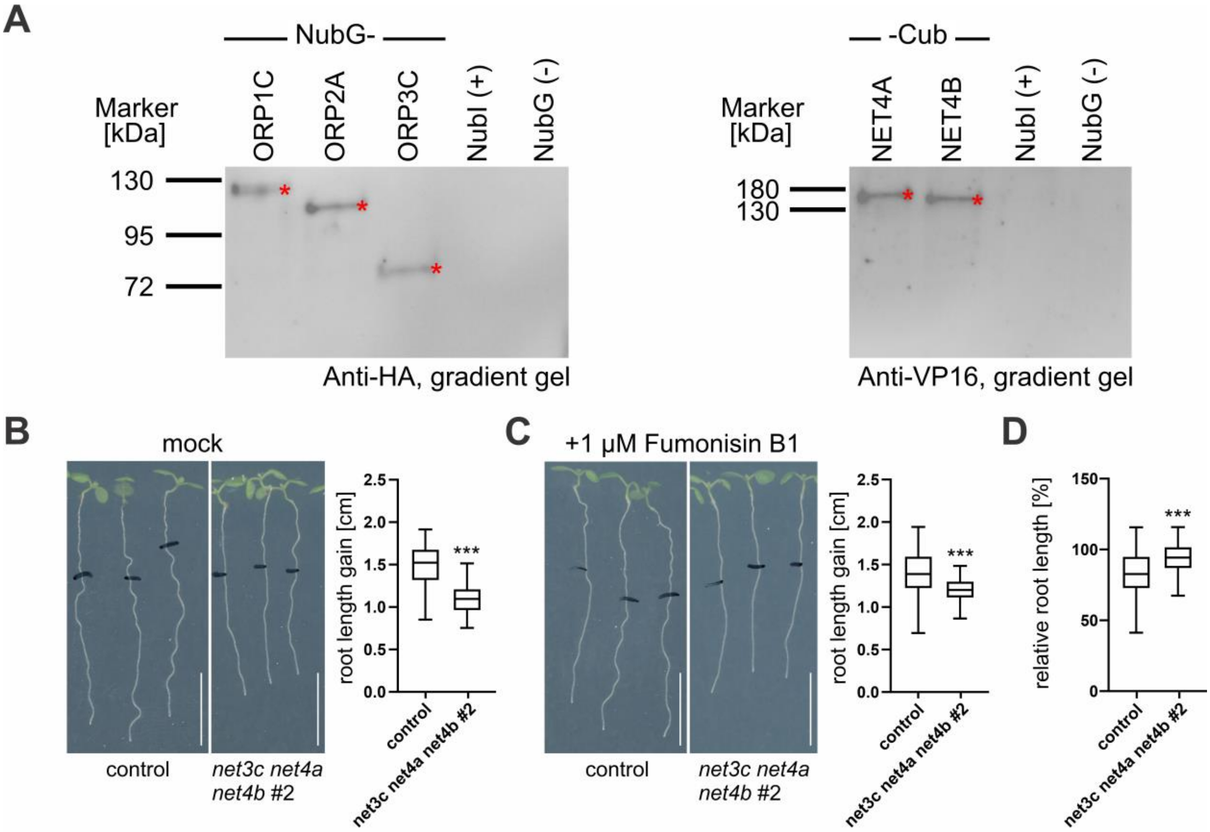


**Fig. S10. Evidence for participation of NET3 and NET4 proteins in lipid homeostasis. (A)** Expression control of mbSUS constructs used in Figure 8. Western blots show detection of NubG-ORP1C, -ORP2A and -ORP3C (left side) as well as NET4A- and NET4B-Cub (right side). Expected size is highlighted by red asterisks **(B)** Arabidopsis root growth (3 days) of Col-0 (n=55) and *net3c net4a net4b* (n=51) upon transfer of 4 days old seedlings from half-strength MS medium onto solvent control plates. Size at the time of transfer is indicated. **(C)** Root length measurements upon transferring 4 days old Col-0 (n=51) and *net3c net4a net4b* (n=47) seedlings onto 1 µM Fumonisin B1 for additional 3 days. **(D)** Relative root growth of Col-0 and *net3c net4a net4b* were calculated (Fumonisin B1 treatment/mock control). Box limits represent 25th-75th percentile, the horizontal line the median and whiskers minimum to maximum values. Significant differences are shown and were analyzed by Student's t-test; ****P* < 0.001. Scale bars = 0.5 cm.
